# The γ-Tubulin Ring Complex promotes mitotic spindle integrity and acts as a microtubule minus-end cap during mitosis

**DOI:** 10.64898/2026.04.10.717779

**Authors:** Reham Aljumaah, Elizabeth A Turcotte, Sanjana Sundararajan, Vasilisa Aksenova, Alexei Arnaoutov, Mary Dasso

## Abstract

γ-TuRC is the primary microtubule (MT) nucleator in eukaryotic cells. Vertebrate γ-TuRC is composed of γ-tubulin and the γ-Tubulin Complex Proteins (GCPs): GCP2, GCP3, GCP4, GCP5, and GCP6. γ-TuRC localizes to MT-Organizing Centers (MTOCs) and promotes MT nucleation by providing a structural template that mirrors the 13-protofilament symmetry of the MT lattice. To understand the contribution of individual γ-TuRC subunits in mitotic spindle dynamics, we endogenously tagged GCP2, GCP4 or GCP6 with an Auxin-Inducible Degron (AID) tag, enabling precise and rapid depletion of each protein. When depletion occurred before mitotic entry, we observed that cells arrested in prometaphase and that loss of any single subunit resulted in displacement of all γ-TuRC components from spindle poles. In cells with preformed spindles, depletion triggered rapid spindle collapse, demonstrating that γ-TuRC remains essential for the maintenance of spindle integrity. Remarkably, depletion of KIF2A, a MT-depolymerizing kinesin, rescued spindle collapse after γ-TuRC loss, suggesting that KIF2A activity contributes to the spindle instability observed in the absence of γ-TuRC. These findings indicate a dual role of γ-TuRC in mitosis: acting not only as a critical MT nucleation factor for initiation of spindle assembly but also as a stabilizing and capping structure at the minus ends of spindle MTs to preserve spindle integrity.

## Introduction

The γ-Tubulin Ring Complex (γ-TuRC) is the primary microtubule (MT) nucleator in eukaryotic cells (Kollman et al., 2011; Oakley, 2023; Zheng et al., 1995). γ-TuRC provides a structural template that mirrors the 13-protofilament symmetry of the MT lattice, thereby promoting MT polymerization by overcoming the energetic barrier to spontaneous nucleation (Kollman et al., 2011; Kraus et al., 2023). γ-TuRC is composed of γ-tubulin and γ-tubulin complex proteins (GCPs) (Murphy et al., 2001; Murphy et al., 1998). In yeast, γ-tubulin associates with spindle pole body proteins Spc97 and Spc98, homologues of GCP2 and GCP3, to form the γ-tubulin small complex (γ-TuSC) (Knop et al., 1997; Kollman et al., 2008). In vertebrates, additional subunits GCP4, GCP5, and GCP6 (each bound to γ-tubulin) are also present in γ-TuRC (Fava et al., 1999; Martin et al., 1998; Moritz et al., 1998; Murphy et al., 1998; Oegema et al., 1999; Zheng et al., 1995). Elegant structural studies have established that vertebrate γ-TuRC contains 14 spokes made of GCP subunits in a distinct stoichiometry and order consisting of 5 copies of GCP2, four to five copies of GCP3, two to three copies of GCP4 and one copy of GCP5 and GCP6 each (Consolati et al., 2020; Liu et al., 2020; Wieczorek et al., 2020; Wurtz et al., 2022). The localization of γ-TuRC to MT-organizing centers (MTOCs), as well as its intrinsic MT nucleation activity are tightly regulated by γ-TuRC interacting proteins such as pericentrin, CEP192, NEDD1 and CDK5RAP2 (Fong et al., 2008; Haren et al., 2006; Haren et al., 2009; Lin et al., 2015; Luders et al., 2006; Manning et al., 2010).

In fungi and *Drosophila*, GCP2 and GCP3 (γ-TuSC subunits) have been reported to be necessary and sufficient for the targeting of γ-Tubulin to centrosomes and for MT nucleation (Anders et al., 2006; Fujita et al., 2002; Lin et al., 2015; Venkatram et al., 2004; Verollet et al., 2006; Xiong and Oakley, 2009). In mitotic human cells, depletion of γ-TuRC using small interfering RNAs (siRNA) causes an increased incidence of monopolar spindles, loss of γ-Tubulin from centrosomes and loss of MT nucleation (Bahtz et al., 2012; Cota et al., 2017; Gomez-Ferreria et al., 2012). However, different studies showed variable requirements for the different γ-TuRC subunits in mitosis (Bahtz et al., 2012; Cota et al., 2017). The interpretation of these studies is limited by the siRNA-mediated depletion methods used for three reasons: First, siRNA depletion frequently produces cell-to-cell variations in the extent of depletion. Second, the extensive time required for siRNA depletion means that cells necessarily entered mitosis without the targeted subunits, so that it was not possible to assess their function in spindle stability after the establishment of a bipolar spindle. Finally, siRNA against specific subunits often resulted in co-depletion of other subunits, thereby making it difficult to dissect the role of individual γ-TuRC components (Bahtz et al., 2012; Cota et al., 2017).

To better understand how γ-TuRC functions during mitosis, we employed a rapid Auxin-Inducible Degradation (AID) strategy to acutely deplete individual γ-TuRC subunits during mitosis. We found that depletion of any single GCP subunit abolished γ-TuRC localization at centrosomes and led to prometaphase arrest when depletion occurred before mitotic entry. Strikingly, the loss of γ-TuRC from completely assembled spindles triggered rapid spindle collapse, revealing that γ-TuRC is required not just for spindle formation but also for the maintenance of spindle integrity. Notably, co-depletion of KIF2A rescued the collapse phenotype, indicating that KIF2A-mediated MT depolymerization contributes to spindle instability in the absence of γ-TuRC. Together, these findings suggest that γ-TuRC not only nucleates spindle MTs but also plays an essential role in stabilizing and capping spindle MT minus ends, thereby preserving spindle architecture after its initial formation.

## Results and Discussion

### Co-dependent localization of γ-TuRC subunits at centrosomes during mitosis

To selectively deplete specific γ-TuRC subunits, we used CRISPR/Cas9 to biallelically tag the endogenous loci of GCP2, GCP4 and GCP6 with 3 copies of the mini-Auxin Inducible Degron (mAID) tag and NeonGreen (NG) fluorescence tag in pseudodiploid DLD-1 colorectal cancer cell lines (Figure 1A and Supplementary Figure 1) as described in our earlier work (Aksenova et al., 2022). To drive the degradation of the targeted proteins, we also engineered the locus encoding the housekeeping protein RCC1 to express the plant protein TIR1 tagged as a fusion with RCC1 and an Infrared Fluorescent Protein (IFP). Once expressed, the TIR1 moiety was cleaved off, to yield RCC1 ^IFP^. RCC1 ^IFP^ served as a marker for chromatin in live imaging experiments, while TIR1 catalyzed ubiquitination and depletion of the target protein after auxin addition (Aksenova et al., 2022). Throughout this report, these cells will be referred to as GCP2 ^NG:AID^, ^AID:NG^ GCP4 and ^AID:NG^ GCP6.

**Figure 1:**
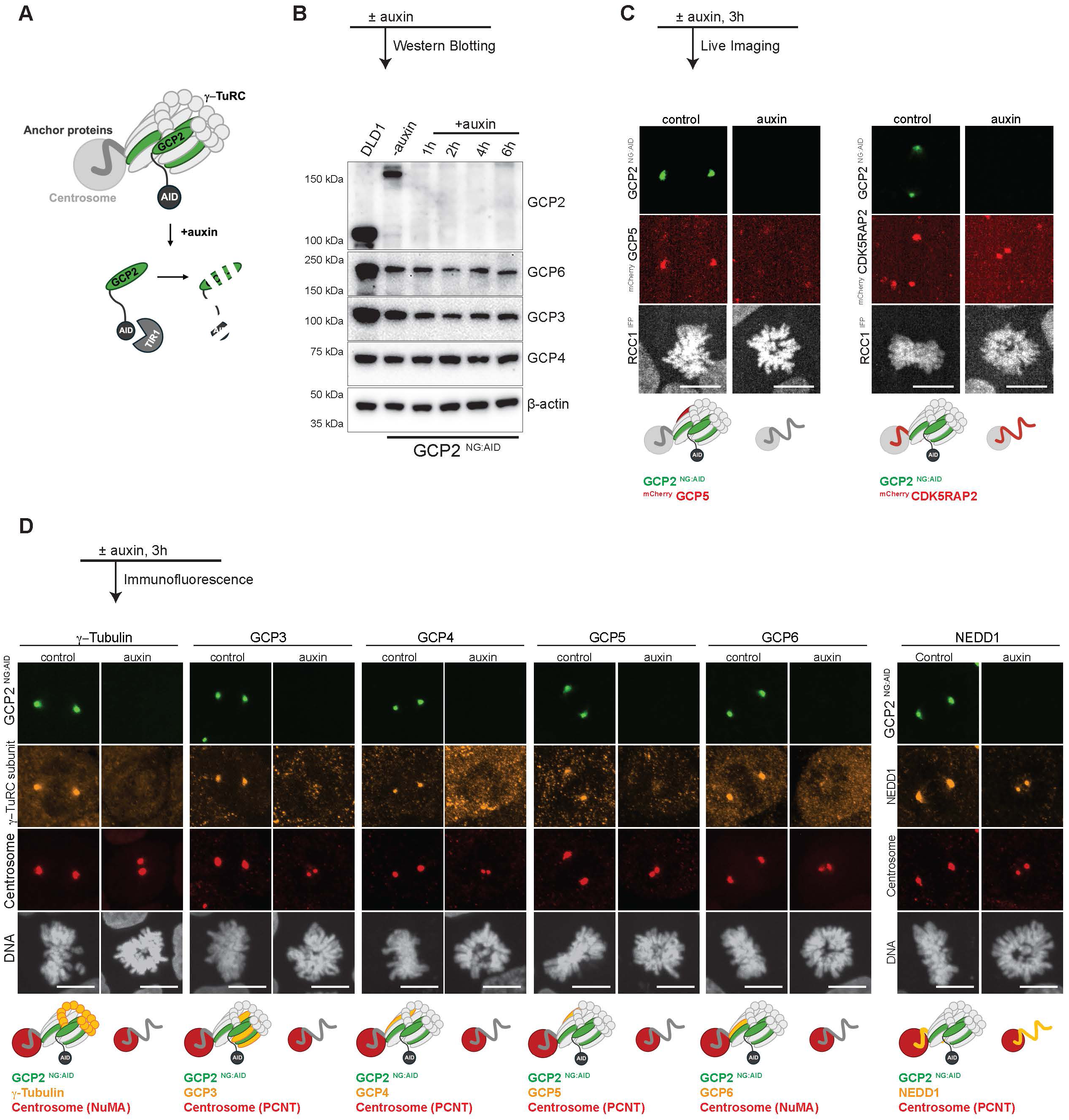
GCP2 - dependent localization of γ-TuRC subunits at centrosomes. A- Schematic showing auxin mediated depletion of GCP2 (protein of interest). GCP2 (green) is tagged with NG for visualization and AID for targeted proteasome mediated depletion upon addition of plant hormone auxin that binds TIR1 to recruit the E3 ligase machinery. B- Western blot showing persistence of γ-TuRC subunits in GCP2 ^NG:AID^ cells upon GCP2 depletion. The lanes represent DLD-1 control (lane1) and GCP2 ^NG:AID^ samples (lanes 2-6) for various time points following auxin addition. β-actin was used as the loading control. C- Live imaging of GCP2 ^NG:AID^ simultaneously tagged with ^mCherry^ GCP5 (left panel, red) or mCherry CDK5RAP2 (right panel, red) after 3h of auxin addition along with their respective controls. RCC1 ^IFP^ was used as the chromatin marker and pseudo colored in grey. D- Immunofluorescence staining of various γ-TuRC subunits (orange) in GCP2 ^NG:AID^ cells following 3h of auxin treatment. PCNT (red) was used as a centrosome marker in all cases except for γ-tubulin and GCP6 staining wherein NuMA (red) was used as indicated in the diagrams. DNA was visualized with Hoechst. Data are representations of at least three independent experiments. Scale bars : 10 µm. NG NeonGreen, AID Auxin Inducible Degron, IFP Infrared Fluorescent Protein.

Previous experiments that utilized siRNA found co-depletion of one or more non-targeted subunits during the extended incubation period required for depletion of individual γ-TuRC subunits (Bahtz et al., 2012; Cota et al., 2017). By contrast, GCP2 ^NG:AID^ showed elimination of GCP2 within 1h of auxin addition without substantial changes in the abundance of any of the other γ-TuRC subunits (Figure 1B). Similarly, the ^AID:NG^ GCP4 and ^AID:NG^ GCP6 lines showed rapid and specific depletion of GCP4 and GCP6, respectively (Supplementary Figure 2A-2C).

**Figure 2:**
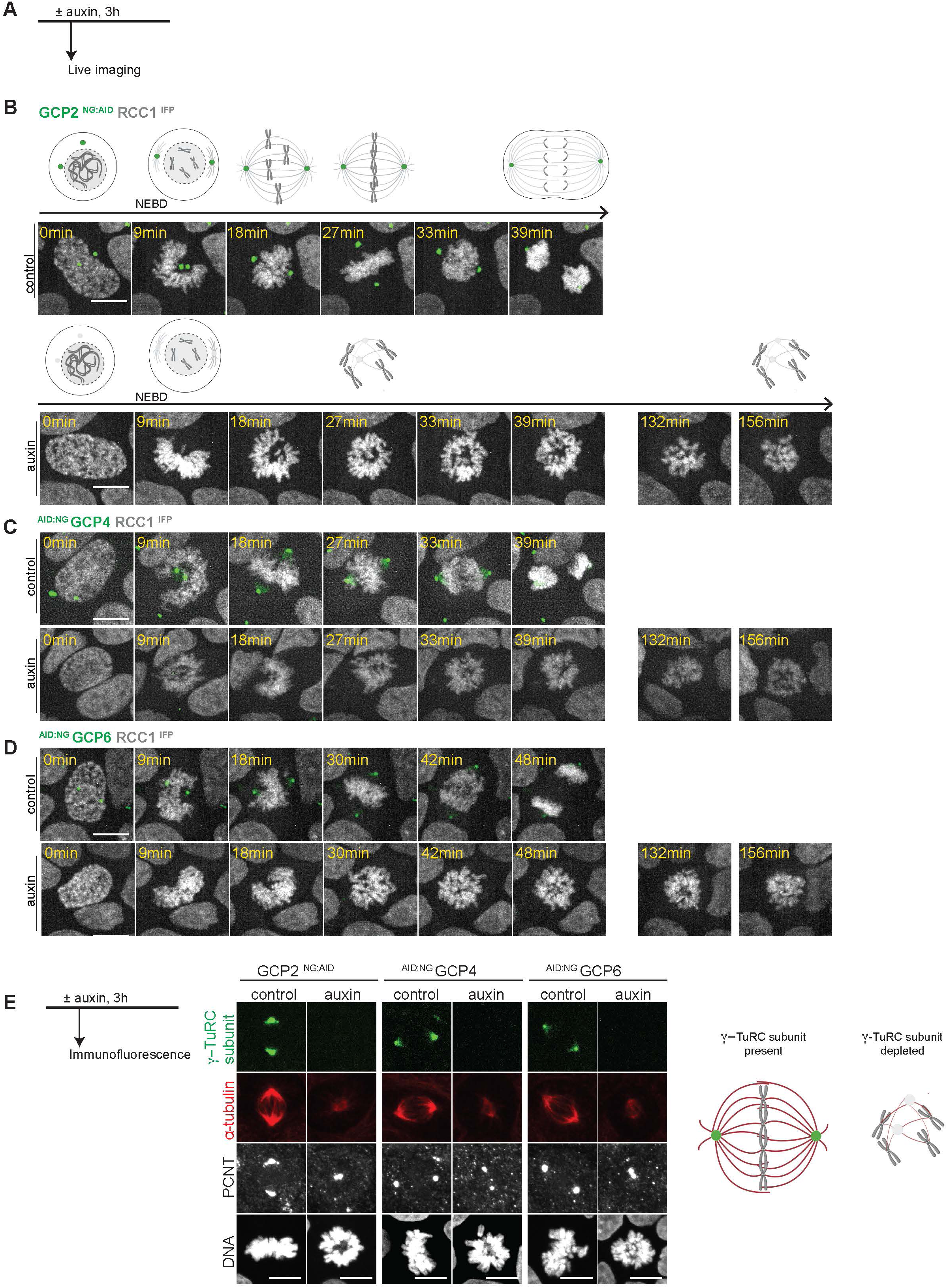
Defective mitotic progression and spindle structures in the absence of γ-TuRC. A- Live imaging protocol for depletion of γ-TuRC subunits prior to mitotic entry. Auxin was added to the culture medium for 3h prior to imaging. B, C, D- Mitotic progression of GCP2 ^NG:AID^, ^AID:NG^ GCP4 and ^AID:NG^ GCP6 cells in control conditions and in the presence of auxin. Note that depletion of any of these subunits resulted in mitotic arrest. Chromatin was visualized using RCC1 ^IFP^. E- Auxin was added for 3h to eliminate GCP2, GCP4 or GCP6 in corresponding AID cell lines. MTs were visualized using α-tubulin (red). γ-TuRC subunits (green) and PCNT (grey) used as the centrosome marker are shown. DNA was visualized using Hoechst. Scale bars : 10 µm.

The persistence of non-targeted subunits after depletion via AID-mediated degradation allowed us to test the co-dependence of γ-TuRC subunits and γ-TuRC recruiting factors for correct localization. To do this, we expressed GCP5 or CDK5RAP2 as fusions with a fluorescent mCherry tag in GCP2 ^NG:AID^ cells and performed live imaging of mitotic cells after the addition of auxin. We observed that GCP2 depletion resulted in the loss of GCP5 from the centrosomes, indicating that its recruitment requires GCP2, while CDK5RAP2 localization was unchanged (Figure 1C). We extended this analysis using immunofluorescent staining of γ-tubulin, GCP3, GCP4, GCP5, GCP6 and NEDD1 (Figure 1D). We observed that in addition to CDK5RAP2, NEDD1 was retained at centrosomes in the absence of GCP2, indicating that the full γ-TuRC complex depends upon GCP2 for correct localization while γ-TuRC recruiting factors do not. Moreover, GCP4 or GCP6 depletion similarly resulted in the loss of localization of all other γ-TuRC subunits from centrosomes without displacement of NEDD1 (Supplementary Figure 2D-F).

Our findings are consistent with earlier reports that NEDD1 localization does not require GCP2 (Manning and Kumar, 2007). Previous reports found differences between γ-TuRC subunits regarding whether they are required for maintenance of others at the centrosome (Bahtz et al., 2012; Cota et al., 2017), but we observed that all γ-TuRC components tested were essential for localization of all others. We suspect that this difference may reflect the greater efficiency for depletion through degron-mediated destruction. Together, our data show that γ-TuRC subunits uniformly depend upon each other for their residence at centrosomes, but that recruiting factors do not require γ-TuRC localization to maintain their centrosome association.

### γ-TuRC is required for establishment of the mitotic spindle apparatus

Depletion of γ-TuRC subunits or γ-TuRC recruiting factors using siRNA-based approaches results in an increased incidence of monopolar spindles (Bahtz et al., 2012; Cota et al., 2017; Gomez-Ferreria et al., 2012). Importantly, the time required for effective depletion using these methods necessitates that the cells would enter mitosis under the depleted conditions. To assess cells entering mitosis after AID-directed depletion of specific γ-TuRC subunits, we performed live imaging of the GCP2 ^NG:AID^, ^AID:NG^ GCP4 and ^AID:NG^ GCP6 cell lines.

While control cells progressed efficiently through mitosis, loss of any of these subunits during interphase resulted in a prolonged arrest in prometaphase (Figure 2A-D), indicating that γ-TuRC is essential for mitotic progression. We further examined spindle morphology in these depleted cells. We found that while control cells displayed typical bi-polar spindles, cells lacking γ-TuRC subunits had fewer, poorly organized MT masses (Figure 2E and Supplementary Figure 3). The morphology of these structures was indicative of a loss of γ-TuRC-dependent MT nucleation and consistent with findings after depletion of γ-TuRC subunits using siRNA-based approaches (Bahtz et al., 2012; Cota et al., 2017; Gomez-Ferreria et al., 2012). Notably, the chromosomes maintained kinetochore (KT) – MT attachments following γ-TuRC depletion (Supplementary Figure 4). Taken together, our data confirm that each of these γ-TuRC subunits is essential for spindle assembly and mitotic progression.

### γ-TuRC is required for maintenance of the mitotic spindle

The rapid depletion of AID-tagged proteins allowed us to further examine whether γ-TuRC is also required to keep the spindle intact once it is established, in a manner that could not be addressed using siRNA-based strategies. To test the role of γ-TuRC in spindle maintenance, we tagged endogenous α-tubulin (TubA3), the plus-end directed MT motor protein Eg5, or the MT minus-end/spindle pole marker NuMA with mCherry to allow live visualization of spindle structures within the GCP2 ^NG:AID^ cell line. These cells were first arrested in metaphase by treatment with ProTAME, a drug that inhibits the Anaphase Promoting Complex (APC), so that they formed complete bipolar spindles. We then added auxin, allowing us to image both the degradation of GCP2 through the loss of NG signal and each of the other tagged proteins via mCherry (Figure 3A).

**Figure 3:**
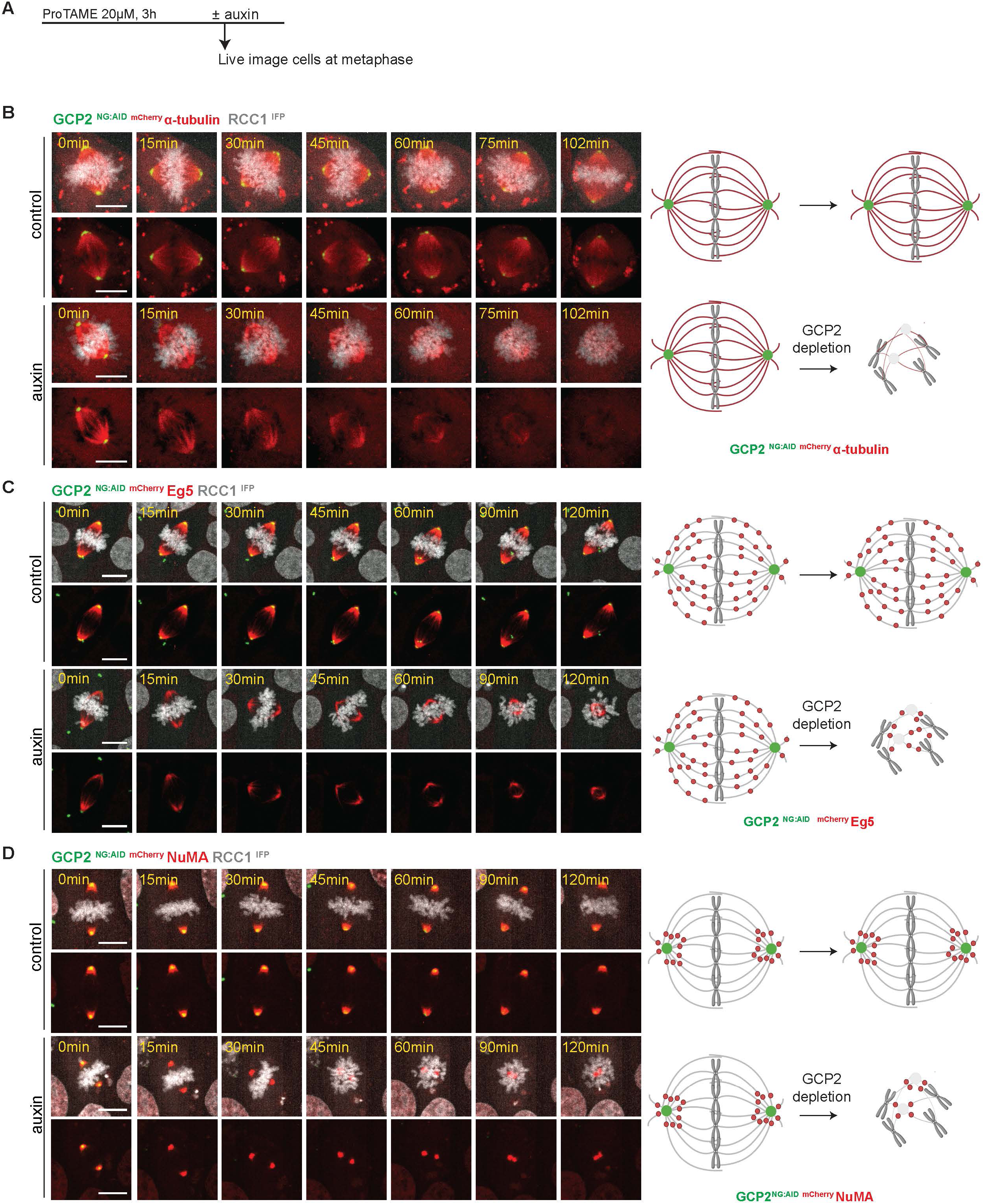
Collapse of assembled spindles upon GCP2 depletion. A- Protocol for live imaging of cells arrested with ProTAME in metaphase prior to GCP2 depletion to assess its role in spindle maintenance. Auxin was added at the time of starting the live imaging analyses. B, C, D- Live imaging analysis of metaphase arrested GCP2 ^NG:AID^ cells co-labeled with either mCherry α-tubulin, ^mCherry^ Eg5 or ^mCherry^ NuMA as indicated. RCC1^IFP^ was used to visualize chromatin. Scale bars : 10 µm.

As GCP2 was depleted, removing γ-TuRC from centrosomes, we observed gradual loss of α-tubulin signal as well as a collapse of bipolar spindles (Figure 3 B). The re-distribution of ^mCherry^ Eg5 and ^mCherry^ NuMA further indicated spindle shortening upon GCP2 depletion (Figure 3C, D). Spindle collapse was observed in metaphase GCP2 ^NG:AID^ cells in the absence of synchronization (Supplementary Figure 5A), arguing that ProTAME treatment did not promote spindle remodeling *per se*, although these experiments were more difficult to conduct because the number of cells that we could observe was limited and because the interpretation of the results might be impacted by mitotic exit. Notably, spindle collapse was also observed in ^AID:NG^ GCP4 and ^AID:NG^ GCP6 cell lines following auxin treatment of ProTAME-synchronized cells, as revealed by immunofluorescent staining for tubulin and the centrosome marker pericentrin. (Supplementary Figure 5B). Together, these observations show that ongoing γ-TuRC activity is essential not only to establish bipolar spindles but also to maintain a bipolar structure after formation.

### γ-TuRC acts as a minus-end MT capping factor

We hypothesized that γ-TuRC association at MT minus ends might serve to cap and stabilize MTs at the centrosome. One prediction of this hypothesis is that γ-TuRC should stably bind to centrosomal MT minus ends. To test this prediction, we arrested GCP2 ^NG:AID^ cells in metaphase with ProTAME and assessed the dynamics of GCP2 using Fluorescence Recovery After Photobleaching (FRAP) experiments by selectively bleaching GCP2 at one centrosome and found no recovery even at 14 min after bleaching (Figure 4A). Similar experiments with ^AID:NG^ GCP4 and ^AID:NG^ GCP6 cells indicated slow or no turnover of GCP4 and GCP6, respectively (Supplementary Figure 6). If this stable population of γ-TuRC served as a minus-end capping structure, we predicted that its removal might render spindle MTs vulnerable to minus-end–directed depolymerizing factors, such as KIF2A, and that reducing their MT depolymerizing activity might counteract the effect of γ-TuRC depletion. To test this idea, we combined auxin-mediated degradation of GCP2 with siRNA-mediated depletion of KIF2A (Supplementary Figure 7).

**Figure 4:**
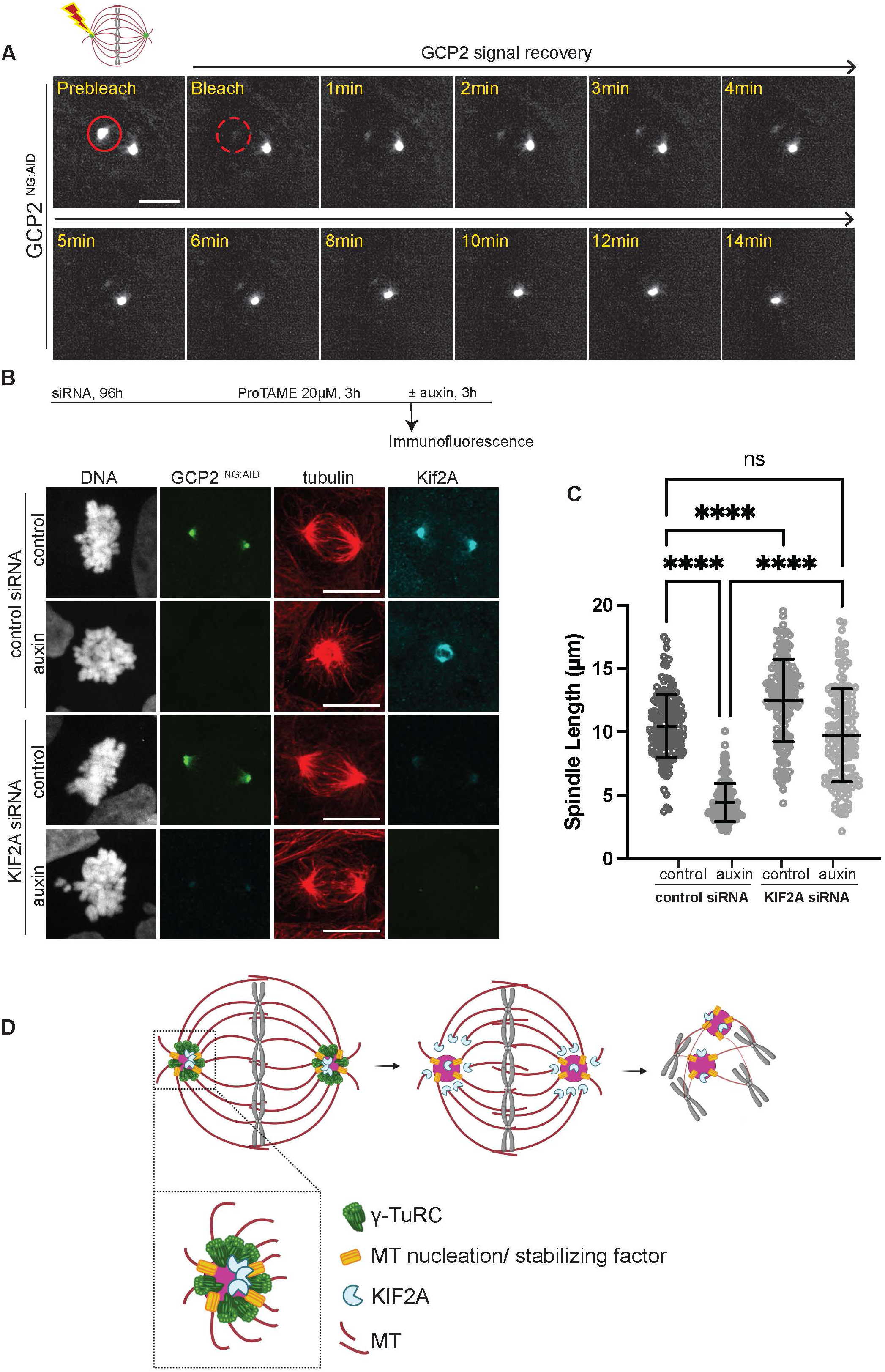
γ-TuRC protects MT minus ends during mitosis. A- FRAP experiment showing slow turnover dynamics for GCP2 at centrosomes. The region at centrosomes targeted for bleaching is indicated by the solid red circle in the prebleach images. Images indicating the region immediately after bleaching and over a course of 14 min are shown. B- (top) Schematic showing experimental protocol for GCP2 depletion coupled with KIF2A siRNA-mediated degradation. GCP2 ^NG:AID^ cells were treated with either control or KIF2A siRNA for 96h followed by ProTAME for 3h and finally auxin for 3h. (bottom) Representative immunofluorescence images showing spindle length. α-tubulin (red, MT) and KIF2A (cyan) staining are shown for the different conditions as indicated. DNA was stained with Hoechst dye. Scale bars : 10 µm. C- Quantification of the total spindle lengths for each condition indicated. Spindle length was measured using tubulin signal as a reference for marking the two ends of the spindle. 150 cells were scored for each condition across three independent experiments. p-values were calculated using one-way ANOVA followed by Tukey post-hoc analysis; ns- p>0.05, ****-p<0.0001. Horizontal bars indicate mean and vertical bars indicate standard deviation. D- Model for the proposed role of γ-TuRC (green) as a minus-end cap that protects MTs (red) from depolymerizing enzyme - KIF2A (cyan). γ-TuRC at the centrosomes functions as the primary MT nucleation and/ or stabilizing factor to aid in the formation and maintenance of a bipolar spindle that facilitates chromosome alignment. Loss of the γ-TuRC exposes the minus ends of γ-TuRC- bound MTs and renders them susceptible to KIF2A activity leading to collapse of the spindle. We propose that a small proportion of MTs are nucleated by γ-TuRC – independent mechanisms (such as the factor depicted in yellow) that keep MTs protected from KIF2A.

Consistent with previous reports, depletion of KIF2A in control GCP2 ^NG:AID^ cells resulted in measurable spindle elongation in the absence of auxin (Figure 4B, C), reflecting its previously established role in regulating spindle length by promoting MT flux through minus-end depolymerization (Bufe et al., 2021). Importantly, co-depletion of KIF2A in auxin-treated cells dramatically rescued spindle collapse after GCP2 depletion (Figure 4B, C), indicating that KIF2A helps to drive spindle collapse in the absence of γ-TuRC. Cumulatively, these findings suggest a model in which γ-TuRC remains stably associated with MT minus ends, capping them throughout mitosis. Loss of γ-TuRC after spindle formation leaves the minus ends susceptible to depolymerizing kinesins such as KIF2A, ultimately leading to spindle collapse (Fig. 4D). This mechanism obviously does not preclude other roles of γ-TuRC in spindle stability or pole focusing (Zeng, 2000).

Interestingly, MT structures persisted after the loss of γ-TuRC from centrosomes, albeit in a highly disorganized fashion (Figure 2E, 3A and Supplementary Figure 5B); also see (Bahtz et al., 2012; Gomez-Ferreria et al., 2012), indicating γ-TuRC–independent MT nucleation mechanisms (Tsuchiya and Goshima, 2021). We speculate that the few MTs that we observed in the absence of γ-TuRC stem from the activity of other MT nucleating / stabilizing factors (Figure 4D). In support of this argument, increasing evidence indicates that liquid–liquid phase separation (LLPS) of selected proteins at centrosomes may generate condensate environments that locally concentrate tubulin dimers and nucleation factors. For instance, in *C. elegans,* such condensates can also recruit key regulators such as TPX2 and ch-TOG, which further stabilize nascent MTs and promote their polymerization, thereby enabling MT assembly even in the absence of canonical γ-TuRC activity (Woodruff et al., 2017). Additionally, other factors such as CAMSAP2 and TPX2, which can initiate MT polymerization and maintain MTs *in vitro* by virtue of phase separation (Brunet et al., 2004; Imasaki et al., 2022; King and Petry, 2020; Roostalu and Surrey, 2017) may contribute to MT maintenance in the absence of γ-TuRC.

## Conclusion

In this study, we show that γ-TuRC plays a critical role in both forming mitotic spindles and in maintaining spindle integrity. Our work also highlights an underappreciated role for γ-TuRC as a minus-end capping complex that prevents MT depolymerization events that would otherwise result in spindle collapse. We demonstrate that γ-TuRC is indispensable for spindle stability throughout mitosis and highlight its essential roles in both MT nucleation and minus-end stability.

## Legends for supplementary figures

**Supplementary Figure 1:**
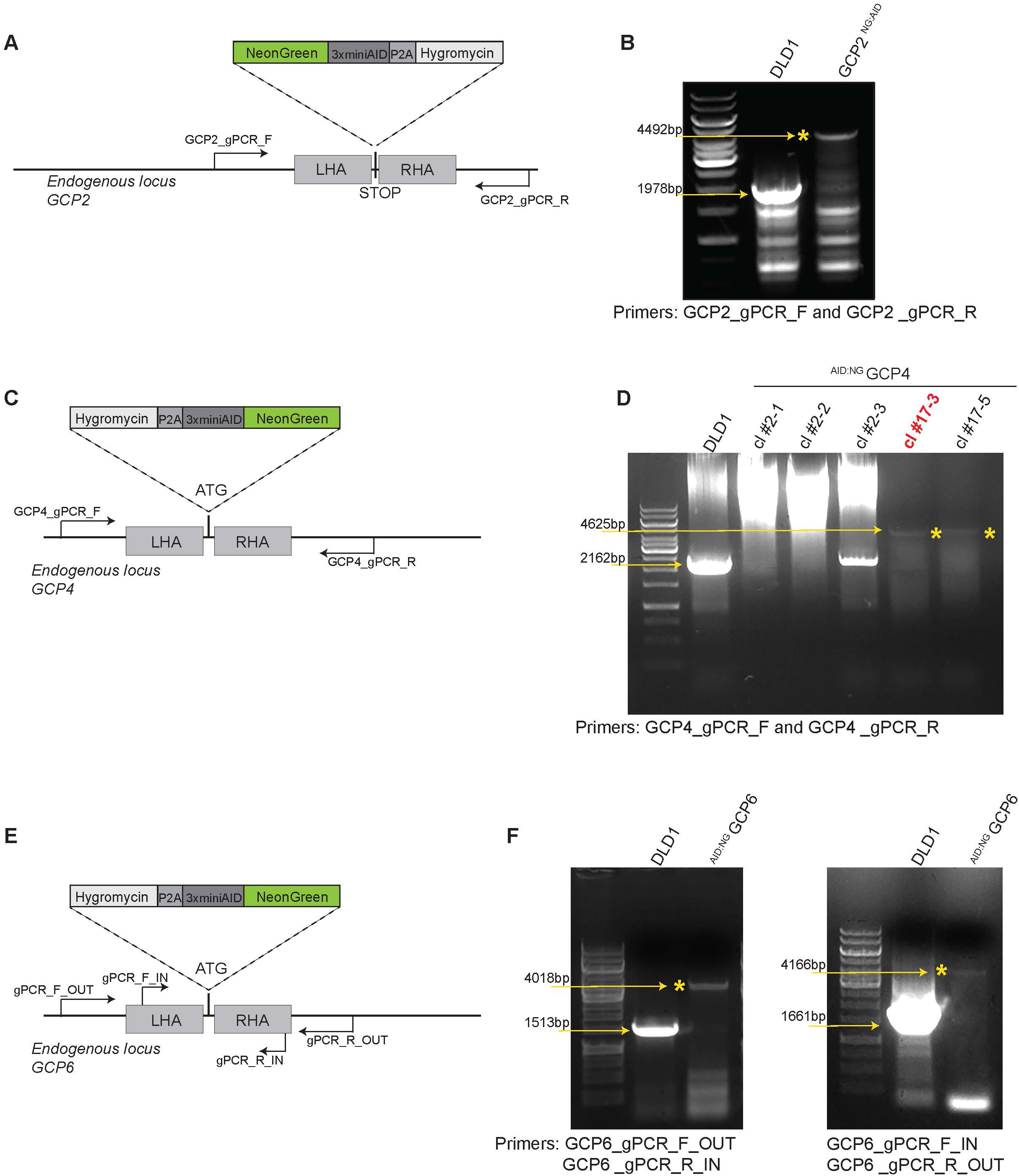
Strategy for tagging individual γ-TuRC subunits and validation of cell lines. A- Schematic for C-terminus tagging of the endogenous *GCP2* locus in DLD-1 cells. GCP2 was tagged with a NG fluorescent tag for visualization and 3X mini-AID tag to mediate targeted depletion. Hygromycin used as a selection marker for clone production was cleaved off by an adjacent P2A site. Primers used for confirming biallelic integration were designed and targeted outside the left and right homology arms as indicated. B- Genomic PCR confirming the biallelic tagging of GCP2 in DLD-1 cells. Control cells (DLD-1) show only the untagged allele (1978 bps) and the GCP2 ^NG:AID^ cells show only the tagged allele (4492 bp, marked by the asterisk). C- Schematic for N-terminus tagging at the endogenous *GCP4* locus with a NG fluorescent tag, a 3X mini-AID tag and a hygromycin selection marker. Primers designed and targeted outside the homology arms were used for confirming the biallelic integration at the GCP4 genomic locus. D- Genomic PCR confirming the biallelic tagging at the *GCP4* locus. Control cells (DLD-1) show only the untagged allele (2162 bps) and the ^AID:NG^ GCP4 cells show only the tagged allele (4625 bp, marked by the asterisk). The clone #17-3 highlighted in red was used in this manuscript. E- Schematic for N-terminus tagging at the endogenous *GCP6* locus with a NG fluorescent tag, a 3X mini-AID tag and a hygromycin selection marker. Primers used for confirming integration were designed and targeted as indicated. F- Genomic PCR confirming the biallelic tagging of GCP6 in DLD-1 cells. (left) The GCP6_gPCR_F_OUT and GCP6_gPCR_R_IN set showed a control band of 1513 bp and 4018 bp band indicating targeted insertion, (right) The GCP6_gPCR_F_IN and GCP6_gPCR_R_OUT combination showed a control band of 1661 bp and 4166 bp band indicating targeted insertion at the *GCP6* locus.

**Supplementary Figure 2:**
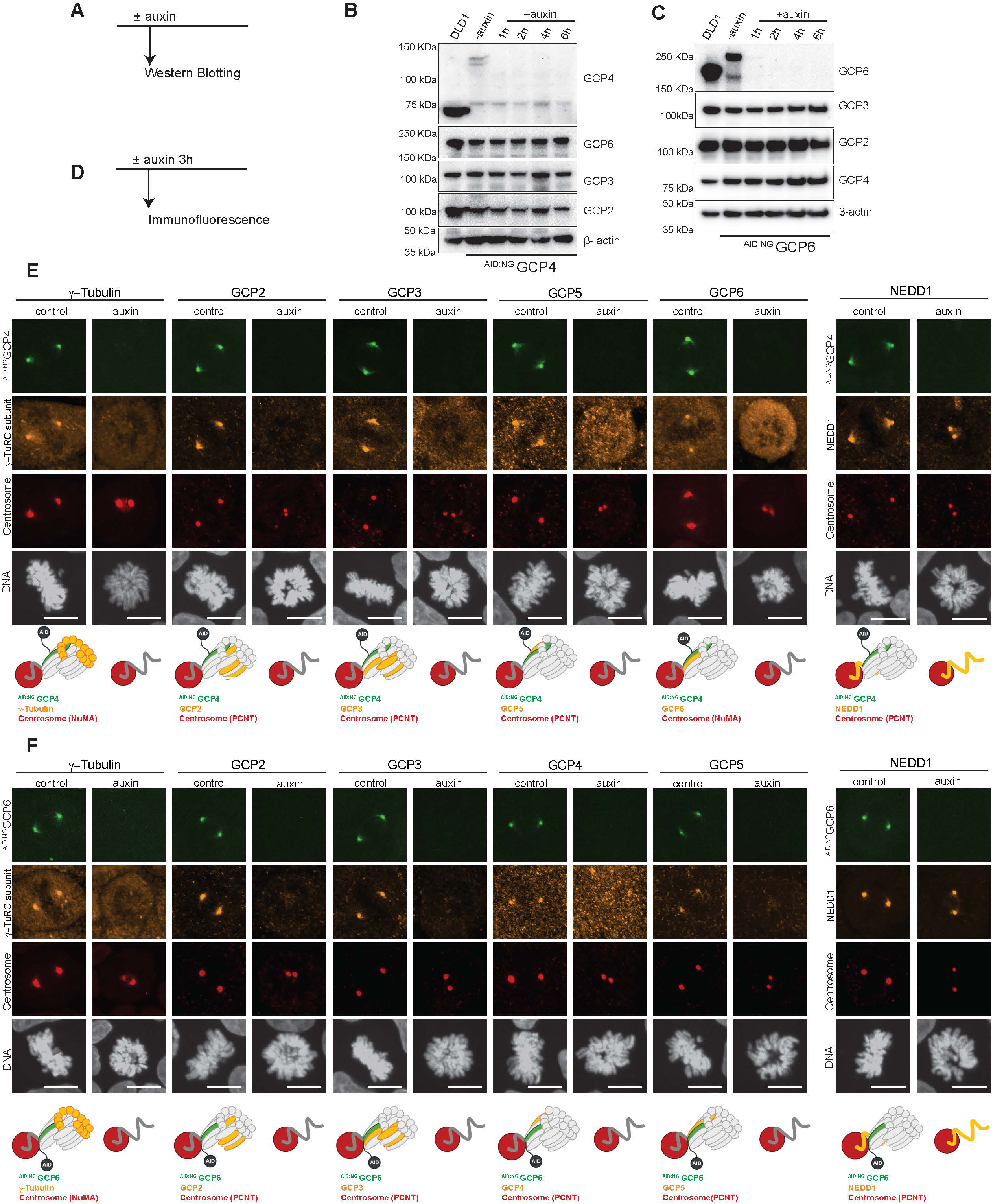
Co-dependence of γ-TuRC subunits for centrosomal targeting. A- Schematic for the protocol used for Western Blotting experiments in B and C. B, C- Western blot indicating persistence of γ-TuRC subunits upon depletion of GCP4 and GCP6 for various time points as indicated. DLD-1 control is depicted in lane 1. β-actin was used as a loading control. D- Schematic for the protocol for Immunofluorescence analysis in E and F. E, F- Immunofluorescence staining for γ-TuRC subunits (orange) upon GCP4 and GCP6 depletion as shown. Centrosomes (red) were marked by PCNT or NuMA as indicated in the diagrams. DNA was visualized with Hoechst. Scale bars : 10 µm.

**Supplementary Figure 3:**
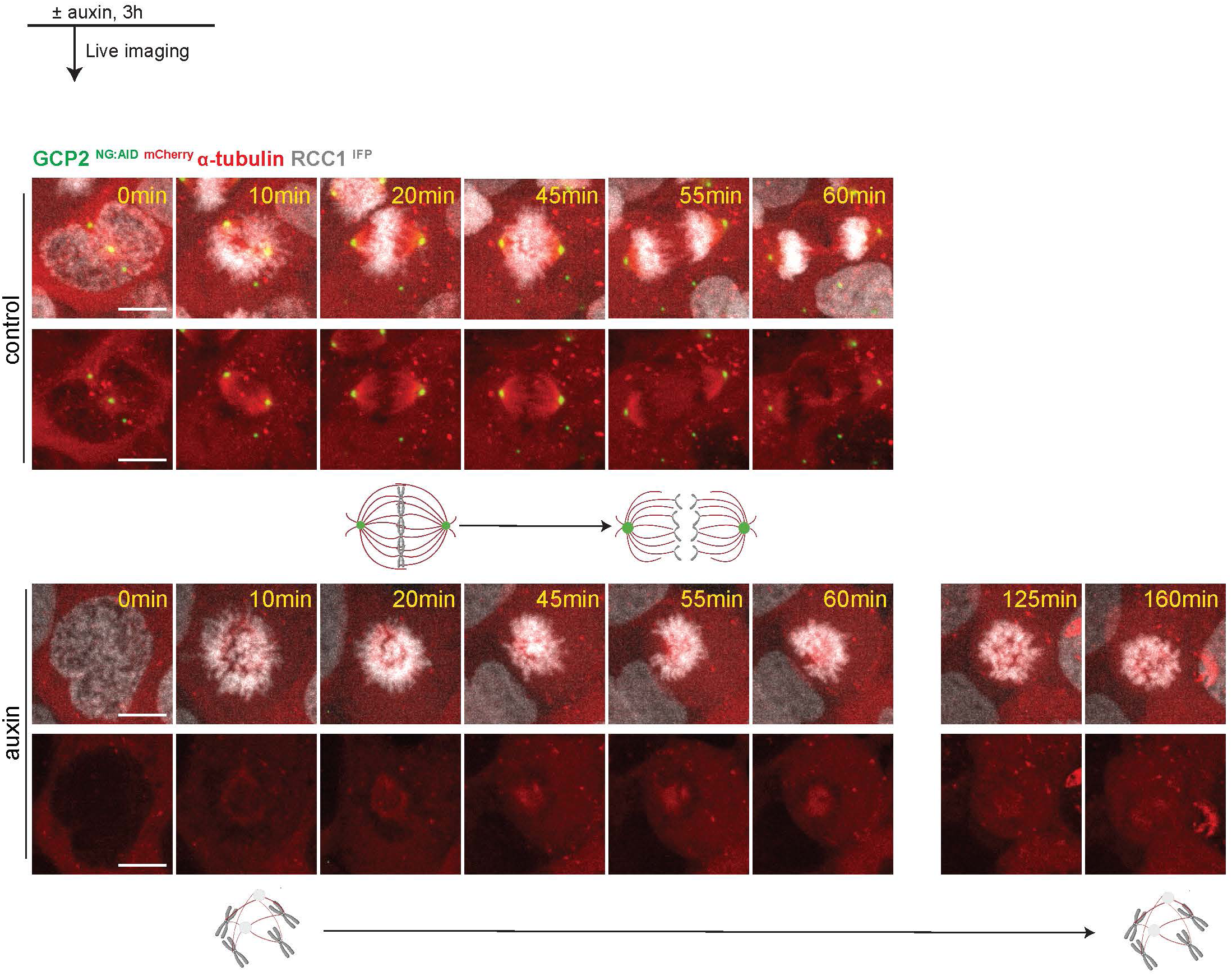
Spindle collapse upon GCP2 depletion visualized by endogenous tubulin tagging. (top) - Schematic for protocol for live imaging experiments to visualize MTs. (bottom) Mitotic progression of GCP2 ^NG:AID^ cells simultaneously tagged with ^mCherry^ α-tubulin. Auxin was added for 3h prior to imaging to mediate GCP2 depletion. RCC1 ^IFP^ was used to visualize chromatin.

**Supplementary Figure 4:**
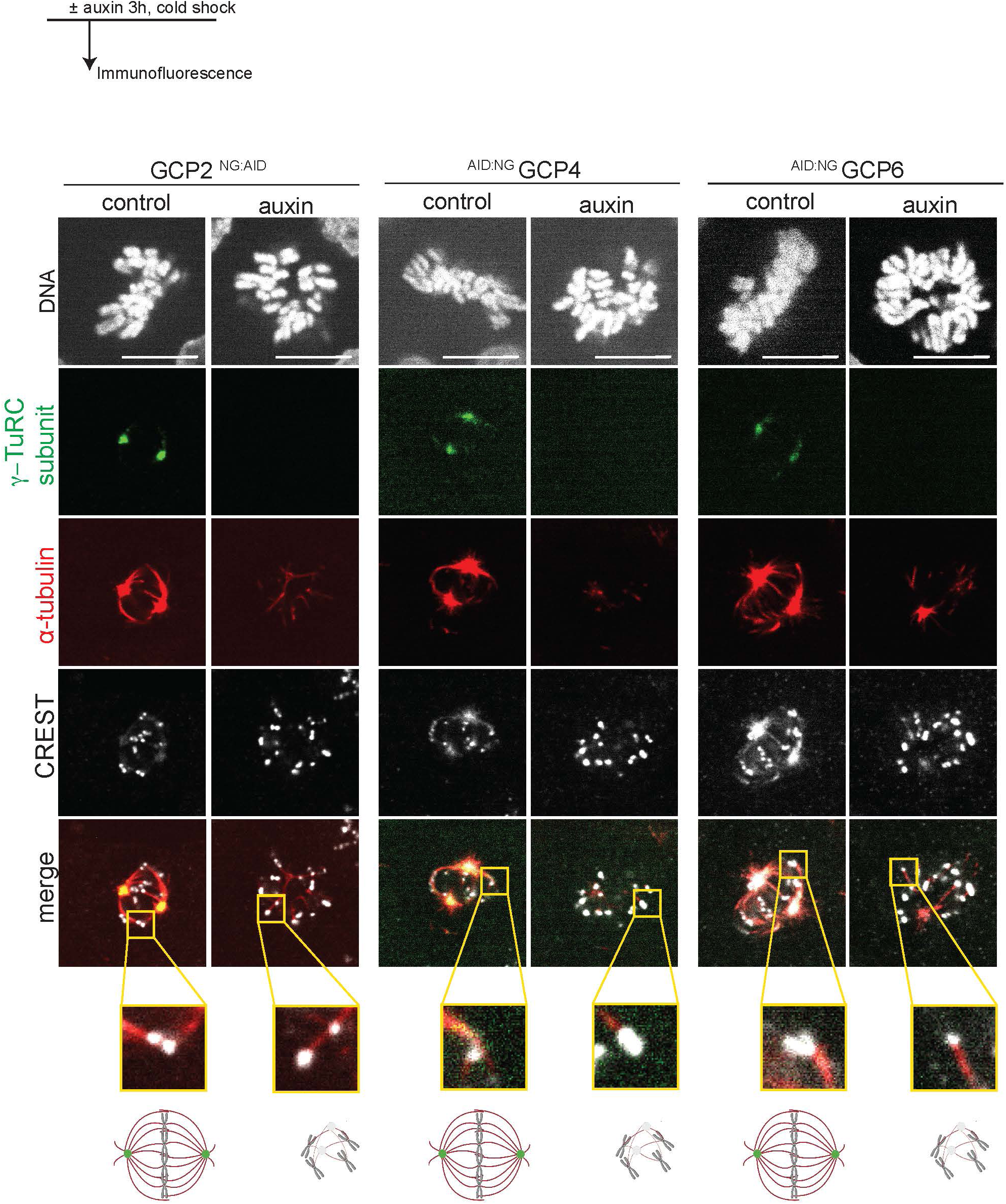
Persistence of KT - MT attachment after spindle collapse induced by loss of γ-TuRC. (top) - Schematic for protocol followed for Immunofluorescence experiment under cold shock for visualizing stable MTs. (bottom)- visualization of cold- stable MTs using α- tubulin (red) upon loss of γ-TuRC subunits as indicated. CREST was used as the KT marker. DNA was visualized with Hoechst. Merged images represent the γ-TuRC subunit indicated (green), α- tubulin (red) and CREST (grey). Images shown are single z sections. Scale bars : 10 µm

**Supplementary Figure 5:**
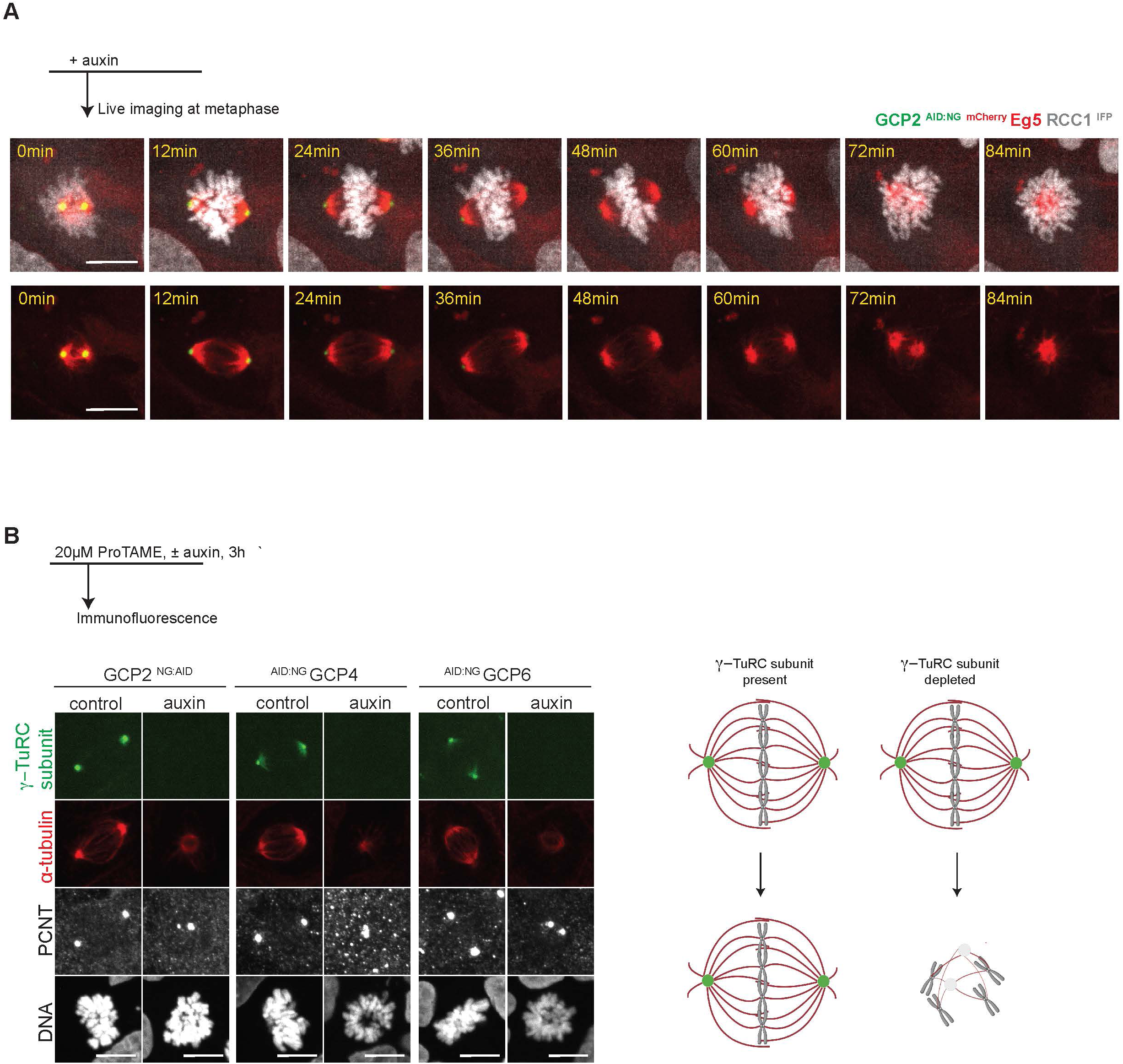
Depletion of γ-TuRC subunits at metaphase results in a weaker collapsed spindle. A- (top) Schematic showing protocol used for live imaging. A cell captured at metaphase was chosen and auxin was added immediately before imaging. (bottom) Mitotic progression of GCP2^NG:AID^ cells simultaneously tagged with ^mCherry^ Eg5 following auxin addition. RCC1 ^IFP^ was used to visualize chromatin. B- (top) Schematic showing protocol used for immunofluorescence staining. ProTAME was used to arrest the cells in prometaphase. Auxin was added 3h prior to fixation of cells to deplete the γ-TuRC subunits as indicated. (bottom) Immunofluorescence staining of metaphase arrested cells for α-tubulin (red). γ-TuRC (green) and PCNT (grey) which was used as the centrosome marker are shown. DNA was visualized with Hoechst. Scale bars : 10 µm

**Supplementary Figure 6:**
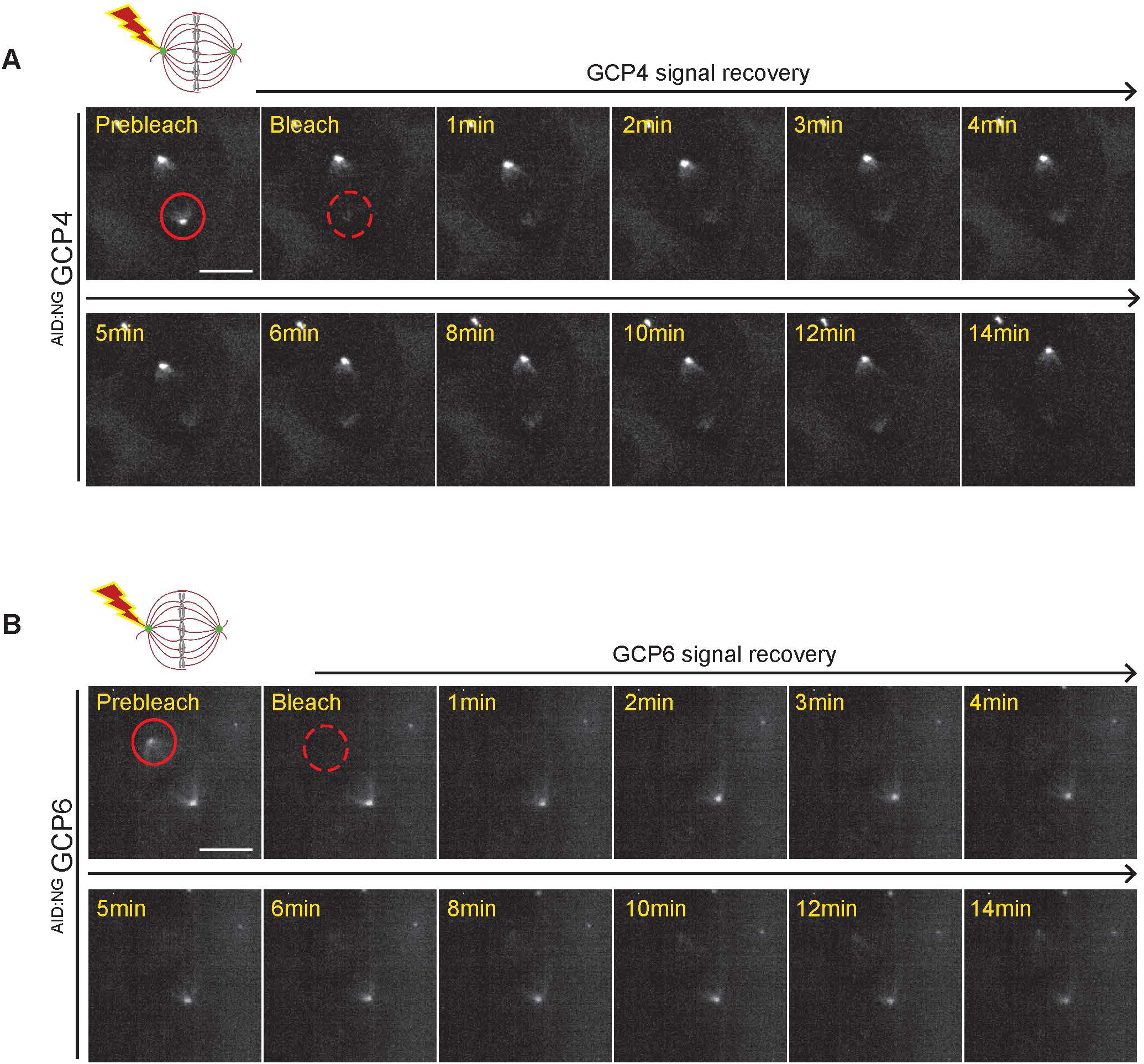
γ-TuRC subunits show slow turnover rate at centrosomes. A, B- FRAP experiment showing GCP4 or GCP6 turnover dynamics. The region at centrosomes targeted for bleaching is indicated by the solid red circle in the prebleach images. Images indicating the region immediately after bleaching and over a course of 14 min are shown. Scale bars : 10 µm.

**Supplementary Figure 7:**
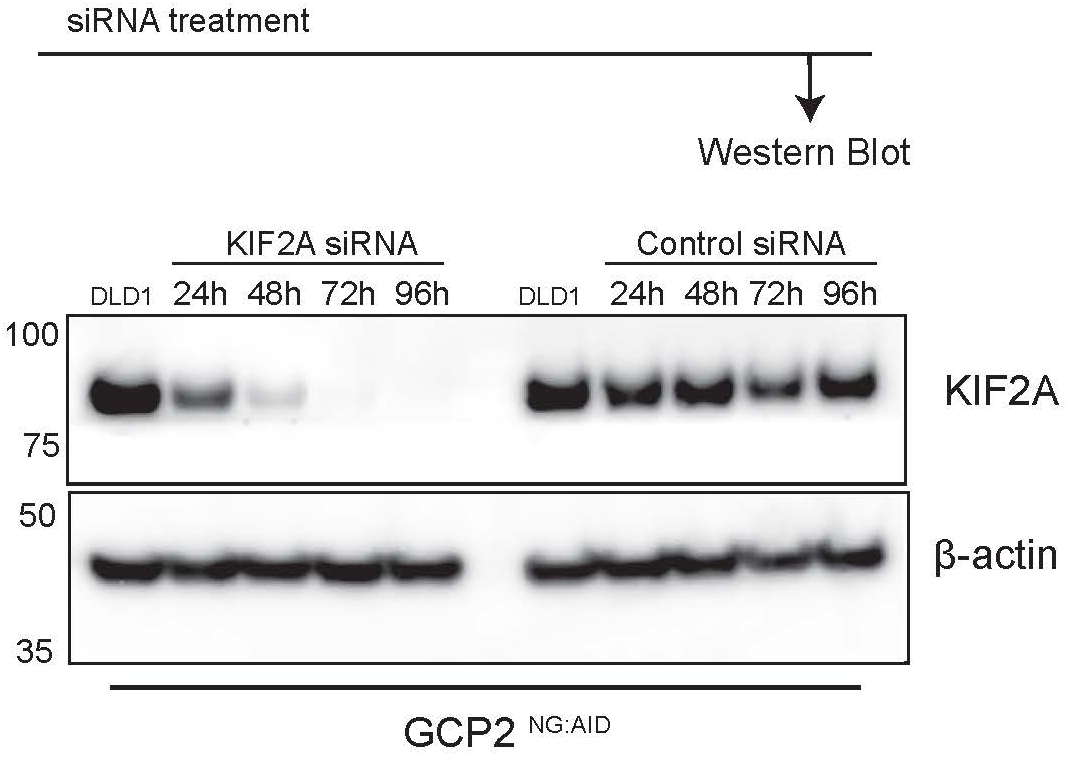
Verification of siRNA mediated KIF2A knockdown. (top) Schematic showing the protocol used for western blots following siRNA mediated depletion. (bottom) Western blot showing confirmation for KIF2A downregulation using siRNA at different timepoints (24h, 48h, 72h, 96h). KIF2A levels are indicated. β- actin was used as a loading control.

## Methods

### Generation of DNA constructs and gene targeting strategies

The CRISPR/Cas9 system was used to endogenously target GCP2, GCP4, GCP6, and RCC1 with NG and 3xminiAID degron or TIR1 using protocol described previously (Aksenova et al., 2022; Aksenova et al., 2020). The GCP2 ^NG:AID^ cell line was further edited to endogenously tag GCP5, CDK5RAP2, tubulin, Eg5 or NuMA with mCherry at the N-terminus. Briefly, the homology arms for generating the donor plasmids were amplified from the genomic DNA of the human DLD-1 cell line. The sequences of hygromycin and TIR1 were obtained from pcDNA-EGFP-AID-BubR1 (Addgene #47330) (Holland et al., 2012) and pBABE TIR1-9Myc (Addgene #47328) (Holland et al., 2012) plasmids respectively by PCR amplification. The NG fluorescent protein and the sequence for 3 copies of the reduced AID tag (3x ∼mini-AID) 65-132 amino acids (Nishimura et al., 2009) were codon optimized and synthesized as described (Aksenova et al., 2020). The cDNAs of mCherry and puromycin resistance gene were amplified from pmCherry-N1 (632523, Clonetech) and pICE vector (Addgene #46960) (Britton et al., 2013) respectively as previously described (Aksenova et al., 2020). The gRNAs sequence was synthesized (Azenta) and integrated into pX330 vector (Addgene #42230) using the protocol from the Zhang Lab (Cong et al., 2013). To visualize α-tubulin, *TUBA1A (TUB3)* locus was targeted with mCherry at N-terminus. The primers used to amplify genomic DNA and for gRNA cloning are listed in Supplementary Table 1. The γ-TURC components were first tagged with NG and AID as indicated in Supplementary Figure 1. The resultant cell lines were then tagged with TIR1 to perform auxin-mediated depletion. To knock-in TIR1, the RCC1(NC_000001.11) housekeeping gene locus was used. CRISPR/Cas9 mediated targeting was performed using a plasmid construct bearing an IFP, 9Myc-TIR1 (Addgene #47328) (Holland et al., 2012) and blasticidin resistance gene (bsr, pOCXIB #631516, Clontech) in such a way that each moiety was separated from the other by a P2A self-cleavage peptide sequence as previously described (Aksenova et al., 2020).

DNA constructs used for subsequent transfection were isolated using the NucleoSpin kit (Takara Bio, 740588.20) according to manufacturer’s instruction.

### Cell Culture, Transfections and Treatments

The human colorectal cancer cell line DLD-1 was cultured in Dulbecco’s Modified Eagle’s Medium (DMEM; ThermoFisher, 10313-021) supplemented with 10% heat-inactivated fetal bovine serum (R&D system Biotechne, S11550), 100 IU/ml penicillin and 100 μg/ml streptomycin (ThermoFisher, 15140-122), and 2 mM GlutaMAX (ThermoFisher, 35050-061) in a 5% CO_2_ atmosphere at 37°C.

To target γ-TURC components with NG and AID or mCherry, cells were plated in density 0.1 x 10^6^ cells/well in 12-well plates a day before transfection. On the day of transfection, complete media was replaced with OptiMEM (Thermofisher, 31985070) without antibiotics two hours before the transfection. Cells on plates were transfected with the donor and guide plasmids (1000 ng total) in a 1:1 proportion using Viafect (Promega, E4981) reagent according to manufacturer’s instructions. After transfection (72 hrs), cells were seeded on 10-cm dishes with the selective antibiotics hygromycin 200 μg/ml (ThermoFisher, 10687010) to select AID-NG-tagged cells, or blasticidin 10 μg/ml to select TIR1 positive clones (ThermoFisher, A1113903), or puromycin 3 μg/ml (ThermoFisher, A11138-03) to select mCherry-tagged cells, until cells formed clones on a plate. Clones with proper localization of the tagged protein and the correct molecular weight on Western blot analysis were further cultured in regular media without selective antibiotics for downstream analysis.

ProTAME, an anaphase-promoting complex/cyclosome inhibitor (Tocris Bioscience, 1362911-19-0), was used to arrest cells at metaphase at a concentration of 20 μM for 2 h prior to the start of the experiment. For protein depletion using the AID system, cells were treated with 1 mM auxin (Sigma, I5148) for the specified period as shown in the schematics for the experiment.

### Genotyping

Genotyping of all cell lines was performed as previously described (Sundararajan et al., 2023). Briefly, genomic DNA isolation was performed with lysis buffer (100 mM Tris-HCl pH 8.0, 200 mM NaCl, 5 mM EDTA, 1% SDS and 0.6 mg/ml proteinase-K (Viagen, 501-pK). This was followed up with ethanol precipitation and resuspension in an elution buffer containing RNAseA (New England Biolabs, Τ3018L). To confirm integration of the construct in the correct locus, the genomic DNA samples were subjected to PCR with primers indicated in the Supplementary Table 2, using GoTaq (Promega, M7123). Information regarding sets of primers used in each case is also provided in Supplementary Figure 1.

### Preparation of Whole Cell Lysates and Western Blotting

DLD-1 cells grown no longer than 10 passages were plated on 6-well plates. When the cells were ∼80% confluent, they were treated with 1 mM auxin for the various timepoints assayed. Cells were washed three times using 1X PBS (Quality Biological, 114-058-101) and pelleted in 1.5 mL eppendorf tubes. The pellet was subsequently lysed using 1X SDS sample buffer. The samples were then boiled, vortexed and ultracentrifuged at 500,000 g (TLA-120.1 rotor) for 10 min at 16°C.

The proteins in the samples were separated by SDS-PAGE performed on precast gradient gels 15-well NuPAGE 4-12% Bis-Tris gels (ThermoFisher, NP0323BOX) and subsequently transferred on to methanol activated Immunobilon-B PVDF 0.45 um membrane (Millipore, IPVH304F0) using semidry transfer (TE77XP SemiDryBlotter, Hoefer). Following the transfer, the membranes were blocked in non-fat milk (5% in 1X PBS, Nestle Carnation, Batch #535304A3) prepared in 1X TBS-T (Tris-buffered saline containing 0.1% Tween-20) for 45 min at room temperature. The membranes were incubated with the suitable primary antibody overnight at 4°C and the following day with the secondary antibodies conjucated with HRP (ECL, anti-rabbit HRP (17671142), anti-mouse (18185770)). for 2 hr at 4°C in dilution 1:10000. Protein bands were detected and visualized using SuperSignal West Pico PLUS Chemiluminescent substrate (ThermoFisher, 34578) using FluorChem Imaging System (ProteinSimple).

The primary antibodies used for western blotting included: GCP2 (TUBGCP2, Novus Biologicals, NBP2-21793), GCP3 (C-3, Santa Cruz Biotechnology, sc-373758), GCP4 (D-5, Santa Cruz Biotechnology, sc-271876), GCP6 (H-9, Santa Cruz Biotechnology, sc2374063), KIF2A (Bethyl, A300-914A), β-actin (Cell Signaling Technology, 4970L) and α-tubulin (Millipore, T6199). β-actin and α-tubulin were used as loading controls for the whole cell lysate samples as indicated.

### Live cell Time-Lapse Imaging

DLD-1 cells were plated and grown on #1.5H Glass Coverslip bottom 4-well chambers (Ibidi, 80427). After the indicated treatment, cells were imaged on an Eclipse Ti2 inverted microscope (Nikon) with a confocal scanning unit (Yokogawa CSU-W1) using a CFI60 Plan Apochromat Lambda 60X oil NA=1.4 WD= 130µm (Nikon). The NIS-Elements AR 5.21.03 software (Nikon) was used to control the microscope. During live imaging, DMEM was replaced with FluoroBrite DMEM (ThermoFisher, A18967-01). The microscope was equipped with a chamber that maintains controlled temperature (37°C), CO_2_ levels, and humidity.

NG and mCherry fluorescent signals were excited using 488-nm and 568-nm laser lines, with a maximum of 20% and 40% power applied, respectively. Optical sections, with z=1µm, were captured every 3 min (Figure 2, Figure 3 and Supplementary Figure 4) and 10 min (Supplementary Figure 3) over a period of 5 h to 10 h total. The images were processed using ImageJ version 1.54b (NIH), Adobe Photoshop (version 27.2.0) and Adobe Illustrator (version 30.1) and the results are presented as maximum intensity projections of the full z-stacks.

### Immunofluorescence staining

DLD-1 cells were plated on 8-well glass-bottom chambers (LAB-TEK II 8-well CV Glass sterile slide system, #) and grown for at least 24 hr before their treatment with 1 mM auxin for 3 hr. The cells were then washed using 1X PBS and fixed and permeabilized using 1X PEM containing 4% paraformaldehyde (PFA, Electron Microscopy Sciences, 15710) and 0.2% TritonX-100 for 15 min. Following this, the cells were blocked with Horse serum (10% in 1X TBS-T, Vector Laboratories, S-2000-20) 1 h at room temperature. Localization of centrosomal proteins and tubulin was visualized using specific primary antibodies and secondary antibodies conjugated with Alexa Fluor (Invitrogen, 568 goat anti rabbit (A11036) 568 donkey anti-mouse 680 (A10037) 680 goat anti mouse (A21058), 647 goat anti-rabbit (A21245), 647 goat anti-human (A21445)). DNA was visualized using 1.5 μg/ml Hoechst 33342 (ThermoFisher, H3569) in all cases as mentioned. Coverslips were subsequently mounted in ProLong Gold antifade reagent (ThermoFisher, P36930).

The primary antibodies used for western blotting included: γ-tubulin (abcam, AB11317), GCP2 (abcam, ab140225), GCP3 (C-3, Santa Cruz Biotechnology, sc-373758), GCP4 (Novus, NBP2-16628), GCP5 (E-1, Santa Cruz Biotechnology, sc-365837), GCP6 (Novus, NBP3–30520), NEDD1 (H-3, Santa Cruz Biotechnology, sc398733), pericentrin (abcam, AB4448), NuMA (F-11, Santa Cruz Biotechnology, sc-365532), KIF2A (Bethyl, A300-914A), and α-tubulin (Millipore, T6199). For staining shown in Figure 2E and Supplementary Figure 5, we diluted the α-tubulin 1:20,000 times to ensure that signal is not saturated.

Images were acquired using Eclipse Ti2 inverted microscope (Nikon) with a confocal scanning unit (Yokogawa CSU-W1) with the samples mounted on a Apochromat TIRF 100X Oil DIC N2 NA = 1.49 WD = 120 µm or a CFI60 Plan Apochromat Lambda 60X oil NA=1.4 WD= 130µm (Nikon). The NIS-Elements AR 5.21.03 software (Nikon) was used to set imaging parameters. Spacing between the z-planes was 0.5µm (Figure 1D, Supplementary Figure 2E, F), 0.3 µm (Figure 2E, Supplementary Figure 4B) and 0.25µm (Figure 4C) . Images were then processed using ImageJ version1.54b (NIH) to uniformly apply brightness and contrast settings to all images. Further processing of the images was using Adobe Photoshop (version 27.2.0) and Adobe Illustrator (version 30.1). All representative images are maximum projections of all the z-slices images.

### Cold-stable MT assay

DLD-1 cells were plated on 8-well glass-bottom chambers (LAB-TEK II 8-well CV Glass sterile slide system, Thermo Scientific, 155409) and grown for at least 24 hr before their treatment with 1mM auxin for 3 hr. The chambers were placed in an ice bath for 10 min to eliminate unstably bound MTs. The cells were then fixed as mentioned above and stained with α-tubulin (Millipore, T6199) and CREST (a gift from Dr. Bill R. Brinkley (Baylor College of Medicine, Houston, TX). Representative images are single z sections.

### siRNA treatment

DLD-1 cells were plated in 12-well plates (containing degreased coverslips when needed for immunofluorescence) and grown for at least 24 hr prior to transfection with siRNA. Degreasing of coverslips was performed with the solution contained 1/2 of 10x Tris/Glycine/SDS buffer (BioRad, 161-0732), 1/4 of ethanol, and ¼ of 1 N NaOH for 5 min to eliminate residual oils. Later, coverslips were washed with water for 15 min, 100% EtOH for 5 min, and dried.

Transfection of control siRNA or KIF2A siRNA (Dharmacon, J-004959-08-0002) was performed using Lipofectamine RNAiMAX reagent (Thermo Fisher, #13778-150) using the manufacturer’s instructions. Following incubation for 24, 48, 72 or 96 hr, the samples were subjected to either western blotting or immunofluorescence. Coverslips that were harvested for immunofluorescence were also subjected to 20µM ProTAME treatment for 2 hr and then 1 mM auxin treatment for 3 hr in addition to the siRNA treatments.

### Measurement of Spindle length

Spindle length was measured using tubulin signal as reference using ImageJ software. The x and y coordinates of the two ends of the spindle were noted. The spindle length was calculated using the formula : spindle length = sq rt ((x_1_-x_2_)^2^ + (y_1_-y_2_)^2^ + (z_1_-z_2_)^2^) where (z_1_-z_2_) = 0.25µm (Chinen et al., 2015).

### Fluorescence Recovery After Photobleaching

DLD-1 cells were plated and grown on 4-well glass-bottom chambers (Ibidi, #80427-90) in FluoroBrite DMEM (ThermoFisher, A1896701) supplemented with 10% FBS (R&D system Biotechne, S11550), as well as antibiotics (100 IU/ml penicillin, 100 μg/ml streptomycin). Cells were imaged live using the CFI Apochromat TIRF 60XC oil immersion objective lens (Nikon) on an Eclipse Ti2 inverted microscope (Nikon) equipped with a confocal scanning unit (Yokogawa CSU-W1) with the stage top incubator maintaining 37°C and 5% CO_2_. Imaging parameters were adjusted using the NIS-Elements AR 5.21.03 software (Nikon). The 488 channel was used to record the fluorescence images with excitation set at 20% of the 488 lasers. Z-axis was adjusted to cover the entire centrosome with the step size of 0.5 µm. The images were acquired over a period of 14 min in total with the images being acquired every 1 min.

### Statistical analysis and reproducibility

All immunofluorescence staining experiments, live imaging, FRAP analyses and western blots were performed at least three times. For quantifying the spindle length in Figure 4, we used three independent experiments; 50 spindles were quantified for each condition in each experiment (total 150 spindles quantified for each condition). Statistical analyses were performed using Graph Pad Prism (Version 10.6.0). One-way ANOVA analysis was used to estimate the statistical significance of the differences in spindle lengths.

## Acknowledgments

RA, ET, SS, VA, AA and MD were supported by the Intramural Research Program of the Eunice Kennedy Shriver National Institute of Child Health and Human Development at the National Institutes of Health, USA (Intramural Project #Z01 HD008954). This research was supported by the Intramural Research Program of the National Institutes of Health (NIH). The contributions of the NIH authors are considered works of the United States Government. The findings and conclusions presented in this paper are those of the authors and do not necessarily reflect the views of the NIH or the U.S. Department of Health and Human Services. We are grateful to Dr. Saroj Regmi for his guidance during the creation of cell lines used in this work. Schematic figures in this manuscript were created using BioRender.com under a license purchased by the National Institutes of Health (NIH). Lastly, we appreciate the members of Mary Dasso’s group for their valuable reviews and constructive feedback during the development of the project and manuscript preparation.

## Abbreviations

γ-TuRC: γ-Tubulin Ring Complex
MT: Microtubule
GCPs: γ-Tubulin Complex Proteins
MTOCs: Microtubule Organizing centers
AID: Auxin Inducible Degron
γ-TuSC: γ-Tubulin Small Complex
NG: NeonGreen
IFP: Infrared Fluorescent Protein
KT: Kinetochore
APC: Anaphase Promoting Complex
LLPS: Liquid-Liquid Phase Separation

---

**Supplementary Table 1:**
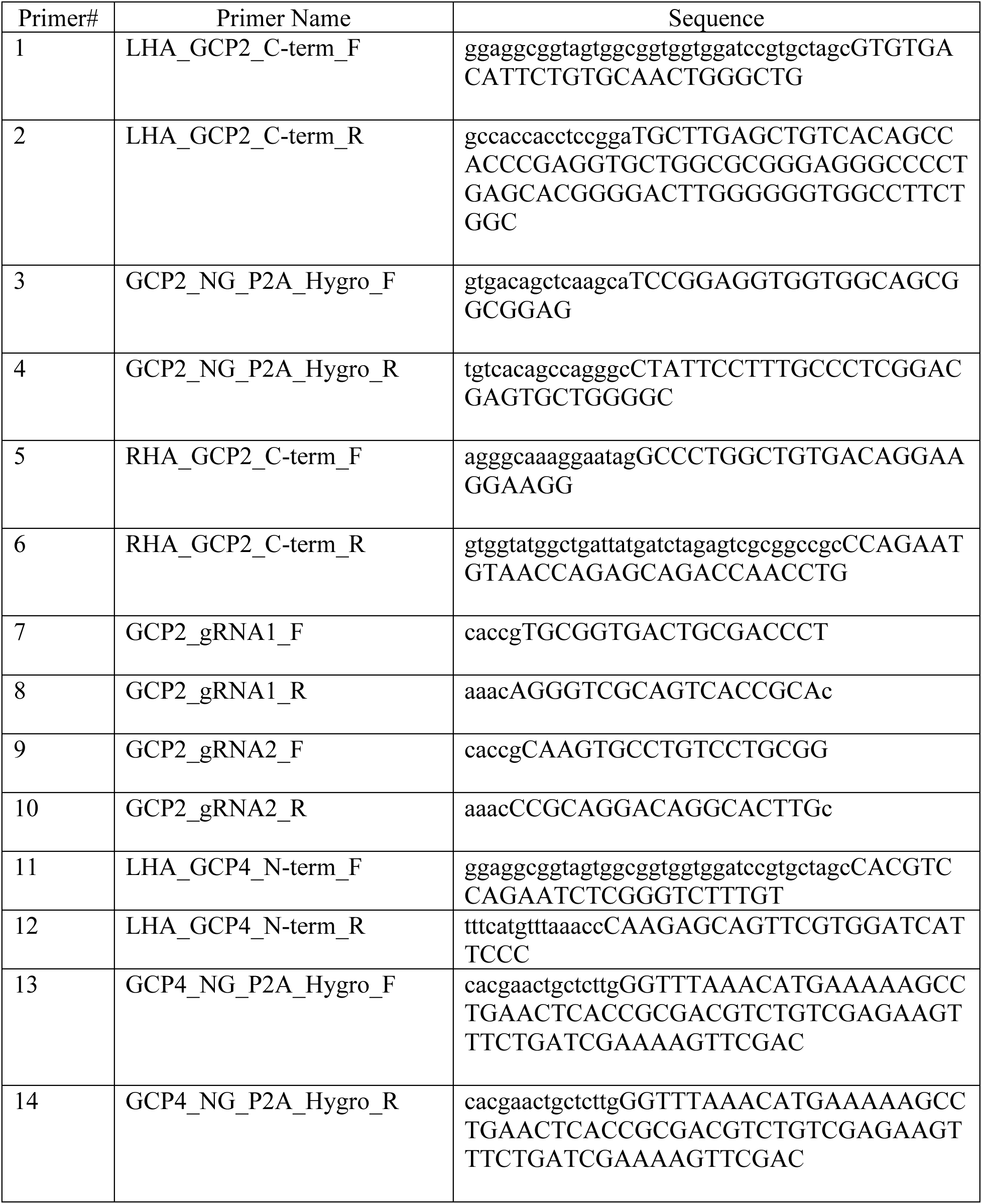
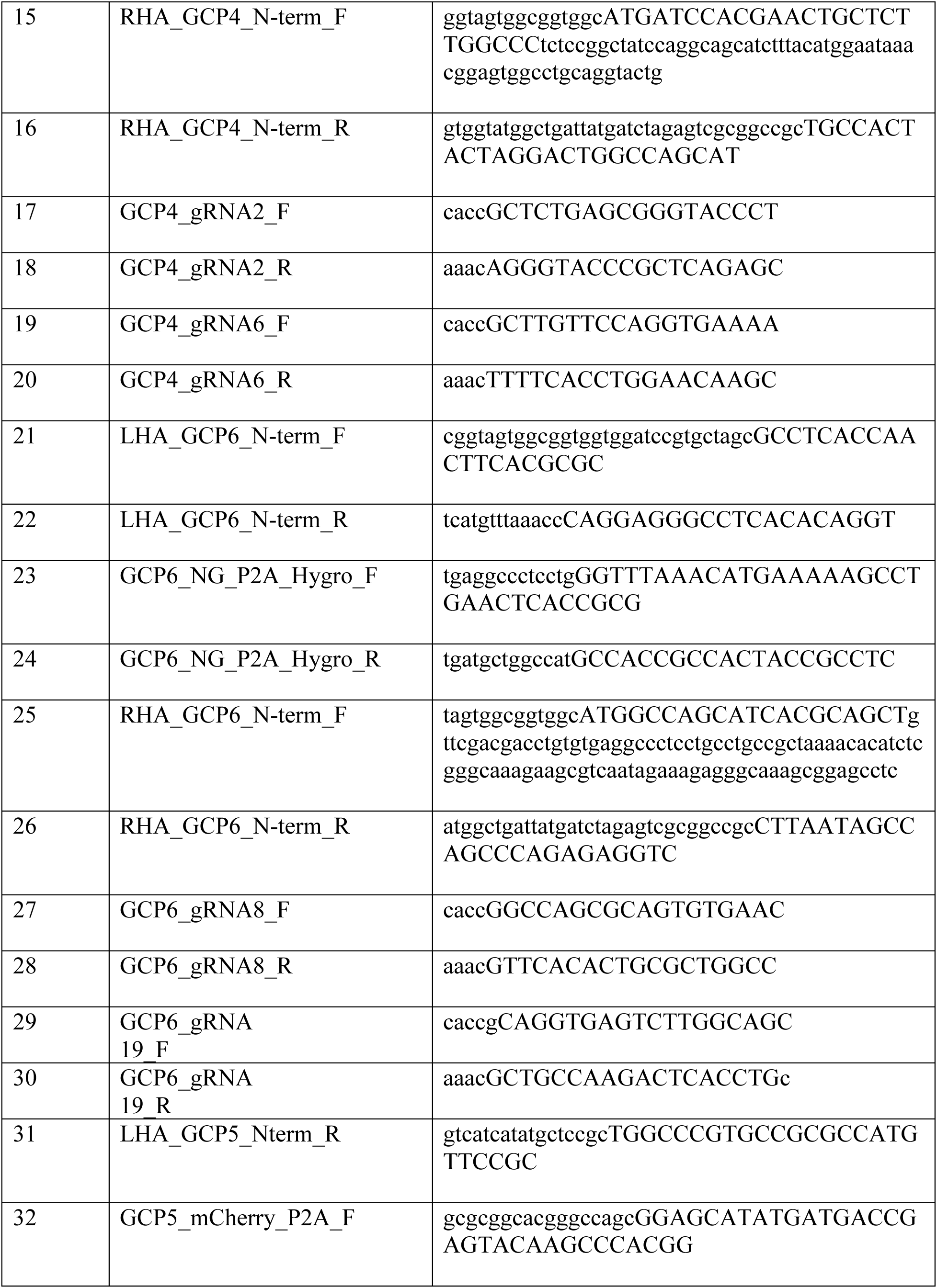
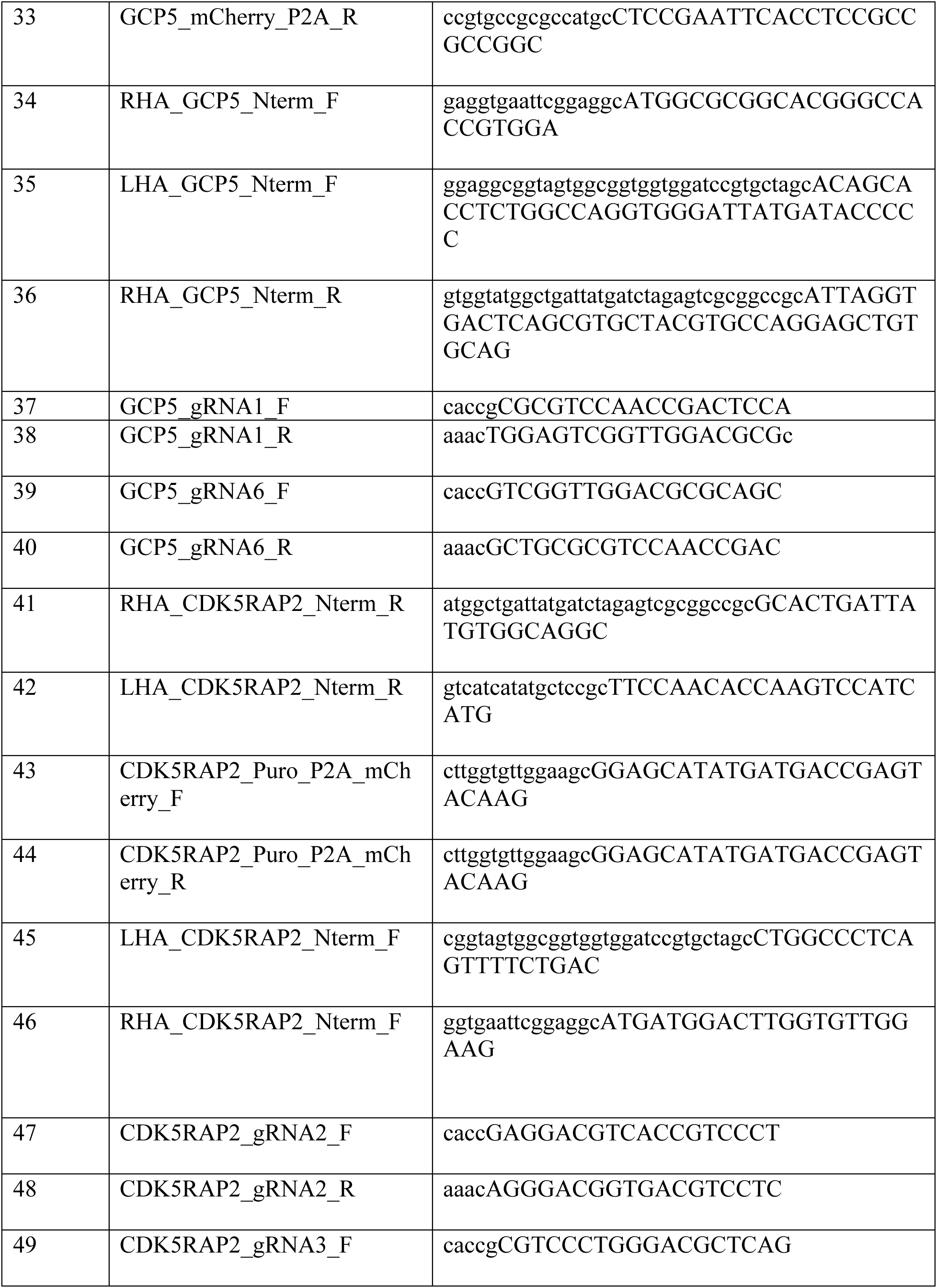
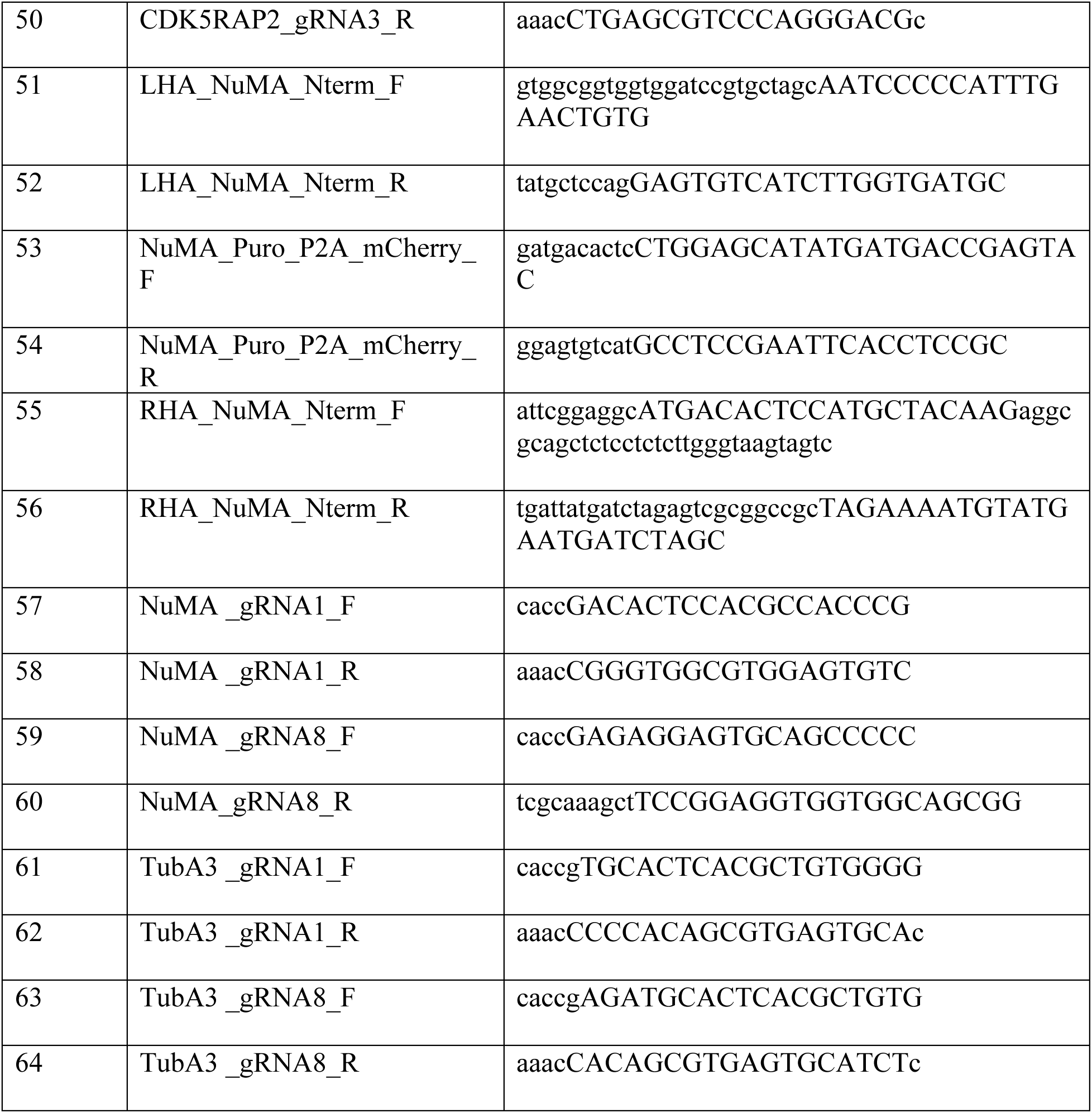
Primers used for the constructing the guide and donor plasmids used in this study.

**Supplementary Table 2:**
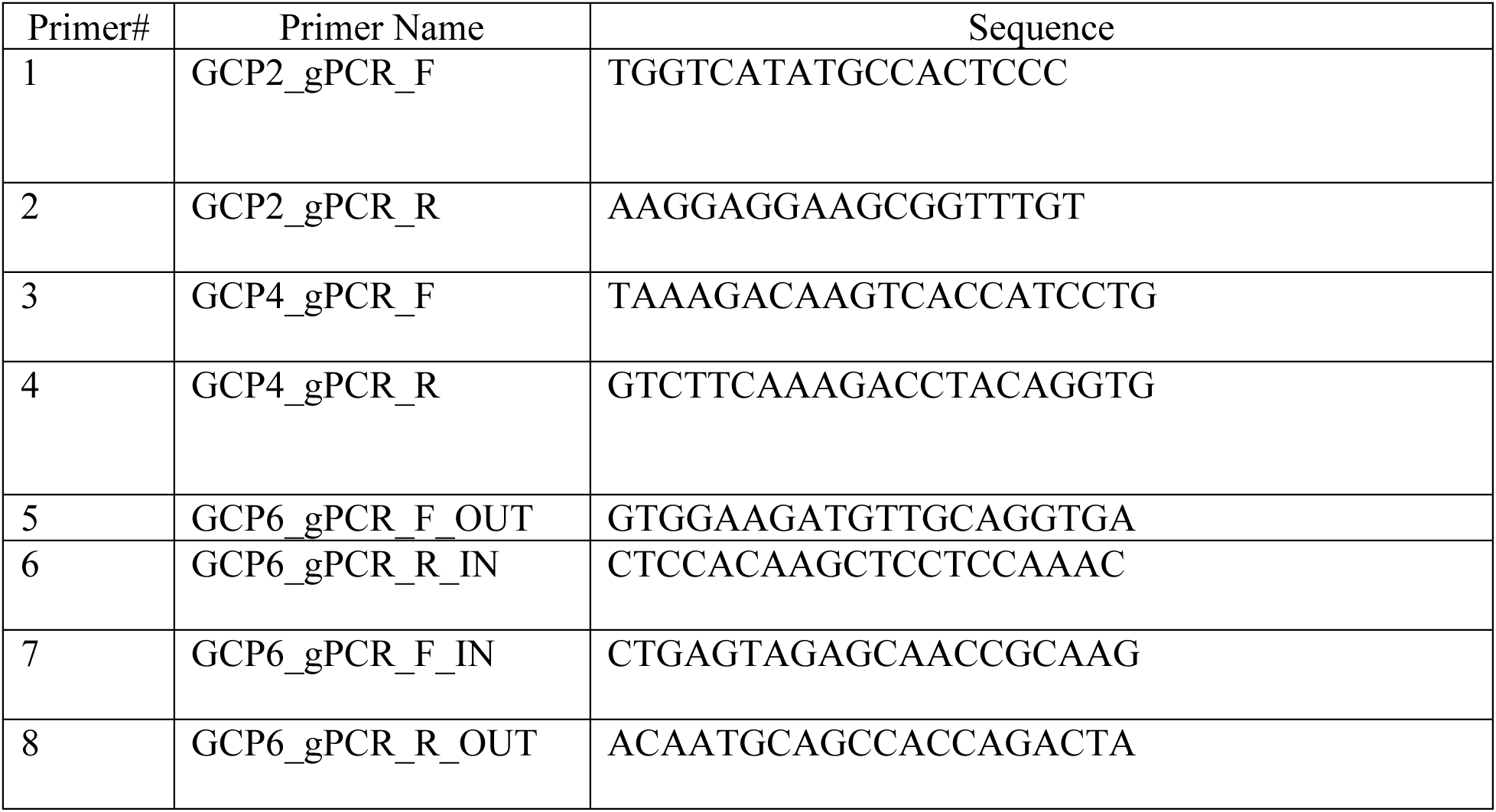
Genomic PCR primers used for the verification of integration in the correct genomic locus.

| Antibody name | Manufacturer | Species | Catalog #, |
| --- | --- | --- | --- |
| GCP2 (anti-TUBGCP2) | Novus Biologicals | Mouse monoclonal | NBP2-21793 |
| GCP2 (anti-TUBGCP2) | abcam | Mouse monoclonal | ab140225 |
| GCP3 (C-3) | Santa Cruz biotechnology | Mouse monoclonal | sc-373758, Lot # B0414 |
| GCP4 (D-5) | Santa Cruz biotechnology | Mouse monoclonal | sc-271876, Lot # G0920 |
| GCP4 (anti-GCP4) | Novus | Rabbit polyclonal | NBP2-16628, Lot #40289 |
| GCP5 (E-1) | Santa Cruz biotechnology | Mouse monoclonal | sc-365837, Lot # C0723 |
| GCP6 (H-9) | Santa Cruz biotechnology | Mouse monoclonal | sc-2374063, Lot #L2624 |
| GCP6 (anti-TUBGCP6) | Novus | Rabbit polyclonal | NBP3-30520, Lot #1 |
| KIF2A (anti-KIF2A) | Bethyl | Rabbit polyclonal | A300-914A |
| α-Tubulin (anti-α-Tubulin) | Millipore | Mouse monoclonal | T6199 |
| β-Actin (beta-Actin (13E5)) | Cell signalling Technology | Rabbit monoclonal | 4970L |
| NEDD1 (H-3) | Santa Cruz biotechnology | Mouse monoclonal | sc-398733 Lot#C0624 |
| Pericentrin | abcam | Rabbit polyclonal | AB4448 |
| γ-Tubulin (gamma-tubulin) | abcam | Rabbit polyclonal | AB11317 |
| NuMA (F-11) | Santa Cruz biotechnology | Mouse monoclonal | sc-365532, Lot #C2211 |

## NIH Intramural Research Program and Employee Publishing Agreement & Manuscript Cover Sheet

By signing this Cover Sheet, the Author, on behalf of NIH, agrees to the provisions set out below, which modify and supersede, solely with respect to NIH, any conflicting provisions that are in the Publisher’s standard copyright agreement (the “Publisher’s Agreement”). If a Publisher’s Agreement is attached, execution of this Cover Sheet constitutes an execution of the Publisher’s Agreement subject to the provisions and conditions of this Cover Sheet.

1. **Indemnification.** No Indemnification or “hold harmless” obligation is provided by either party.
2. **Governing Law.** This agreement will be governed by the law of the court in which a claim is brought.
3. **Copyright.** Author’s contribution to the Work was done as part of the Author’s official duties as an NIH employee and is a Work of the United States Government. Therefore, copyright may not be established in the United States. 17 U.S.C. § 105. If Publisher intends to disseminate the Work outside of the U.S., Publisher may secure copyright to the extent authorized under the domestic laws of the relevant country, subject to a paid-up, nonexclusive, irrevocable worldwide license to the United States in such copyrighted work to reproduce, prepare derivative works, distribute copies to the public and perform publicly and display publicly the work, and to permit others to do so.
4. **No Compensation.** No royalty income or other compensation may be accepted for work done as part of official duties. The author may accept for the agency a limited number of reprints or copies of the publication.
5. **NIH Representations.** NIH represents to the Publisher that the Author is the sole author of the Author’s contribution to the Work and that NIH is the owner of the rights that are the subject of this agreement; that the Work is an original work and has not previously been published in any form anywhere in the world; that to the best of NIH’s knowledge the Work is not a violation of any existing copyright, moral right, database right, or of any right of privacy or other intellectual property, personal, proprietary or statutory right; that where the Author is responsible for obtaining permissions or assisting the Publishers in obtaining permissions for the use of third party material, all relevant permissions and information have been secured; and that the Work contains nothing misleading, obscene, libelous or defamatory or otherwise unlawful. NIH agrees to reasonable instructions or requirements regarding submission procedures or author communications, and reasonable ethics or conflict of interest disclosure requirements unless they conflict with the provisions of this Cover Sheet. Notwithstanding the foregoing, the Author may, consistent with NIH policy, submit a copy of the original work to a public repository.
6. **Disclaimer.** NIH and the Author expressly disclaim any obligation in Publisher’s Agreement that is not consistent with the Author’s official duties or the NIH mission, described at https://www.nih.gov/about-nih. NIH and the Author do not disclaim obligations to comply with a Publisher’s conflict of interest policy so long as, and to the extent that, such policy is consistent with NIH’s own conflict of interest policies.
7. **For Peer-Reviewed Papers to be Submitted to PubMed Central.** The Author is a U.S. government employee who must comply with the NIH Public Access Policy, and the Author or NIH will deposit, or have deposited, in NIH’s PubMed Central archive, an electronic version of the final, peer-reviewed manuscript upon acceptance for publication, to be made publicly available upon the Official Date of Publication, defined in the NIH Public Access Policy as the date on which the Final Published Article is first made available in final, edited form, whether in print or electronic (i.e., online) format. The Author and NIH agree (notwithstanding Paragraph 3 above) to follow the manuscript deposition procedures of the publisher so long as they are consistent with the NIH Public Access Policy, including making the manuscript publicly available without embargo in PubMed Central.
8. **Modifications.** PubMed Central may tag and modify the work consistent with its customary practices and within the meaning and integrity of the underlying work.

The NIH Deputy Director for Intramural Research approves this publishing agreement for NIH staff from the NIH Intramural Program. NIH Institute and Center Directors approve this publishing agreement for NIH staff from their respective Institutes and Centers outside the NIH Intramural Program. The NIH Principal Deputy Director approves this publishing agreement for NIH staff in the Office of the Director. A single, signed copy of this text is maintained for all works published by NIH employees, and contractors and trainees who are working at the NIH. No additional signatures beyond that of the Author are needed.

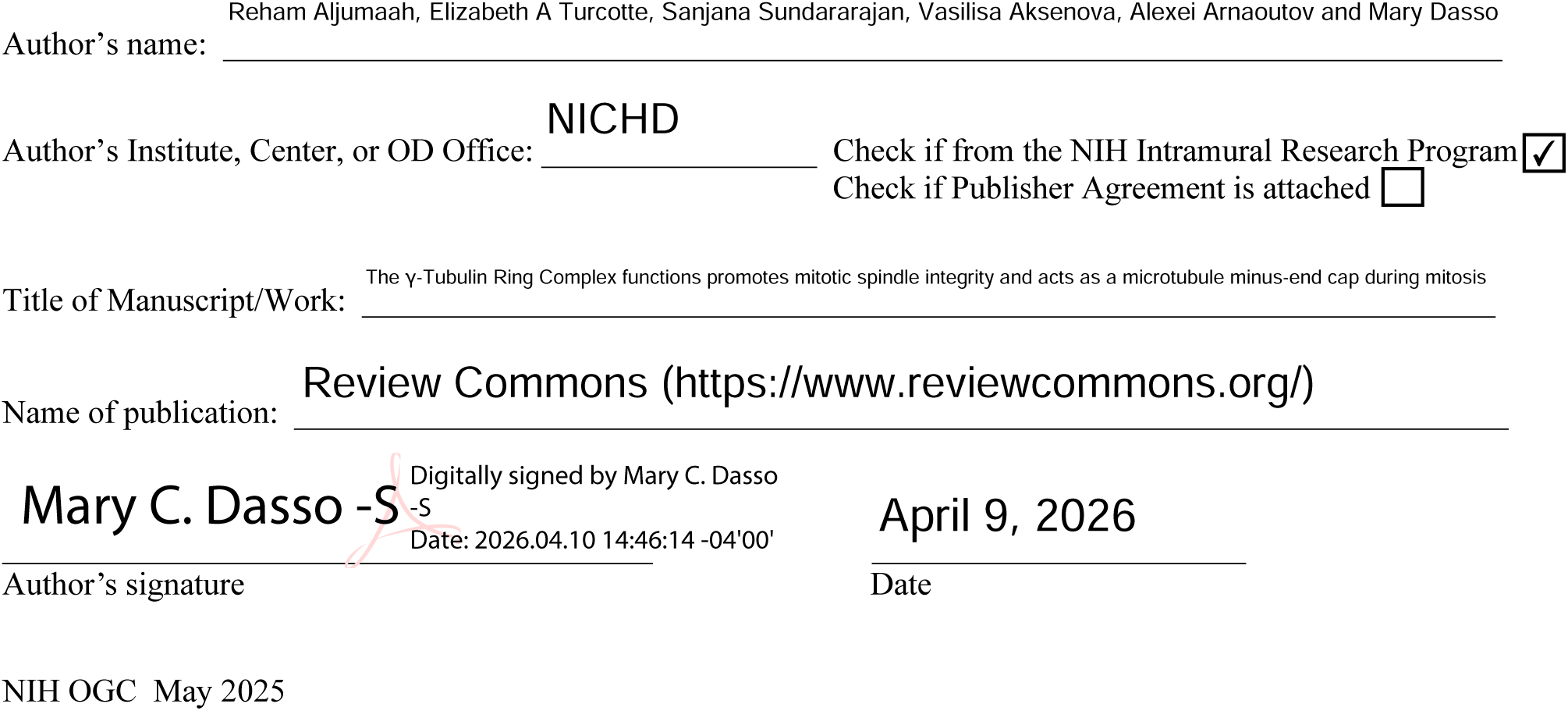

## References

Aksenova, V., A. Arnaoutov, and M. Dasso. 2022. Analysis of Nucleoporin Function Using Inducible Degron Techniques. Methods Mol Biol. 2502:129–150.

Aksenova, V., A. Smith, H. Lee, P. Bhat, C. Esnault, S. Chen, J. Iben, R. Kaufhold, K.C. Yau, C. Echeverria, B. Fontoura, A. Arnaoutov, and M. Dasso. 2020. Nucleoporin TPR is an integral component of the TREX-2 mRNA export pathway. Nat Commun. 11:4577.

Anders, A., P.C. Lourenco, and K.E. Sawin. 2006. Noncore components of the fission yeast gamma-tubulin complex. Mol Biol Cell. 17:5075–5093.

Bahtz, R., J. Seidler, M. Arnold, U. Haselmann-Weiss, C. Antony, W.D. Lehmann, and I. Hoffmann. 2012. GCP6 is a substrate of Plk4 and required for centriole duplication. J Cell Sci. 125:486–496.

Britton, S., J. Coates, and S.P. Jackson. 2013. A new method for high-resolution imaging of Ku foci to decipher mechanisms of DNA double-strand break repair. J Cell Biol. 202:579–595.

Brunet, S., T. Sardon, T. Zimmerman, T. Wittmann, R. Pepperkok, E. Karsenti, and I. Vernos. 2004. Characterization of the TPX2 domains involved in microtubule nucleation and spindle assembly in Xenopus egg extracts. Mol Biol Cell. 15:5318–5328.

Bufe, A., A. Garcia Del Arco, M. Hennecke, A. de Jaime-Soguero, M. Ostermaier, Y.C. Lin, A. Ciprianidis, J. Hattemer, U. Engel, P. Beli, H. Bastians, and S.P. Acebron. 2021. Wnt signaling recruits KIF2A to the spindle to ensure chromosome congression and alignment during mitosis. Proc Natl Acad Sci U S A. 118.

Chinen, T., P. Liu, S. Shioda, J. Pagel, B. Cerikan, T.C. Lin, O. Gruss, Y. Hayashi, H. Takeno, T. Shima, Y. Okada, I. Hayakawa, Y. Hayashi, H. Kigoshi, T. Usui, and E. Schiebel. 2015. The gamma-tubulin-specific inhibitor gatastatin reveals temporal requirements of microtubule nucleation during the cell cycle. Nat Commun. 6:8722.

Cong, L., F.A. Ran, D. Cox, S. Lin, R. Barretto, N. Habib, P.D. Hsu, X. Wu, W. Jiang, L.A. Marraffini, and F. Zhang. 2013. Multiplex genome engineering using CRISPR/Cas systems. Science. 339:819–823.

Consolati, T., J. Locke, J. Roostalu, Z.A. Chen, J. Gannon, J. Asthana, W.M. Lim, F. Martino, M.A. Cvetkovic, J. Rappsilber, A. Costa, and T. Surrey. 2020. Microtubule Nucleation Properties of Single Human gammaTuRCs Explained by Their Cryo-EM Structure. Dev Cell. 53:603–617 e608.

Cota, R.R., N. Teixido-Travesa, A. Ezquerra, S. Eibes, C. Lacasa, J. Roig, and J. Luders. 2017. MZT1 regulates microtubule nucleation by linking gammaTuRC assembly to adapter-mediated targeting and activation. J Cell Sci. 130:406–419.

Fava, F., B. Raynaud-Messina, J. Leung-Tack, L. Mazzolini, M. Li, J.C. Guillemot, D. Cachot, Y. Tollon, P. Ferrara, and M. Wright. 1999. Human 76p: A new member of the gamma-tubulin-associated protein family. J Cell Biol. 147:857–868.

Fong, K.W., Y.K. Choi, J.B. Rattner, and R.Z. Qi. 2008. CDK5RAP2 is a pericentriolar protein that functions in centrosomal attachment of the gamma-tubulin ring complex. Mol Biol Cell. 19:115–125.

Fujita, A., L. Vardy, M.A. Garcia, and T. Toda. 2002. A fourth component of the fission yeast gamma-tubulin complex, Alp16, is required for cytoplasmic microtubule integrity and becomes indispensable when gamma-tubulin function is compromised. Mol Biol Cell. 13:2360–2373.

Gomez-Ferreria, M.A., M. Bashkurov, A.O. Helbig, B. Larsen, T. Pawson, A.C. Gingras, and L. Pelletier. 2012. Novel NEDD1 phosphorylation sites regulate γ-tubulin binding and mitotic spindle assembly. J Cell Sci. 125:3745–3751.

Haren, L., M.H. Remy, I. Bazin, I. Callebaut, M. Wright, and A. Merdes. 2006. NEDD1-dependent recruitment of the gamma-tubulin ring complex to the centrosome is necessary for centriole duplication and spindle assembly. J Cell Biol. 172:505–515.

Haren, L., T. Stearns, and J. Lüders. 2009. Plk1-dependent recruitment of gamma-tubulin complexes to mitotic centrosomes involves multiple PCM components. PLoS One. 4:e5976.

Holland, A.J., D. Fachinetti, J.S. Han, and D.W. Cleveland. 2012. Inducible, reversible system for the rapid and complete degradation of proteins in mammalian cells. Proc Natl Acad Sci U S A. 109:E3350–3357.

Imasaki, T., S. Kikkawa, S. Niwa, Y. Saijo-Hamano, H. Shigematsu, K. Aoyama, K. Mitsuoka, T. Shimizu, M. Aoki, A. Sakamoto, Y. Tomabechi, N. Sakai, M. Shirouzu, S. Taguchi, Y. Yamagishi, T. Setsu, Y. Sakihama, E. Nitta, M. Takeichi, and R. Nitta. 2022. CAMSAP2 organizes a γ-tubulin-independent microtubule nucleation centre through phase separation. Elife. 11.

King, M.R., and S. Petry. 2020. Phase separation of TPX2 enhances and spatially coordinates microtubule nucleation. Nat Commun. 11:270.

Knop, M., G. Pereira, S. Geissler, K. Grein, and E. Schiebel. 1997. The spindle pole body component Spc97p interacts with the gamma-tubulin of Saccharomyces cerevisiae and functions in microtubule organization and spindle pole body duplication. EMBO J. 16:1550–1564.

Kollman, J.M., A. Merdes, L. Mourey, and D.A. Agard. 2011. Microtubule nucleation by γ-tubulin complexes. Nat Rev Mol Cell Biol. 12:709–721.

Kollman, J.M., A. Zelter, E.G. Muller, B. Fox, L.M. Rice, T.N. Davis, and D.A. Agard. 2008. The structure of the gamma-tubulin small complex: implications of its architecture and flexibility for microtubule nucleation. Mol Biol Cell. 19:207–215.

Kraus, J., R. Alfaro-Aco, B. Gouveia, and S. Petry. 2023. Microtubule nucleation for spindle assembly: one molecule at a time. Trends Biochem Sci. 48:761–775.

Lin, T.C., A. Neuner, and E. Schiebel. 2015. Targeting of γ-tubulin complexes to microtubule organizing centers: conservation and divergence. Trends Cell Biol. 25:296–307.

Liu, P., E. Zupa, A. Neuner, A. Böhler, J. Loerke, D. Flemming, T. Ruppert, T. Rudack, C. Peter, C. Spahn, O.J. Gruss, S. Pfeffer, and E. Schiebel. 2020. Insights into the assembly and activation of the microtubule nucleator γ-TuRC. Nature. 578:467–471.

Luders, J., U.K. Patel, and T. Stearns. 2006. GCP-WD is a gamma-tubulin targeting factor required for centrosomal and chromatin-mediated microtubule nucleation. Nat Cell Biol. 8:137–147.

Manning, J., and S. Kumar. 2007. NEDD1: function in microtubule nucleation, spindle assembly and beyond. Int J Biochem Cell Biol. 39:7–11.

Manning, J.A., S. Shalini, J.M. Risk, C.L. Day, and S. Kumar. 2010. A direct interaction with NEDD1 regulates gamma-tubulin recruitment to the centrosome. PLoS One. 5:e9618.

Martin, O.C., R.N. Gunawardane, A. Iwamatsu, and Y. Zheng. 1998. Xgrip109: a gamma tubulin-associated protein with an essential role in gamma tubulin ring complex (gammaTuRC) assembly and centrosome function. J Cell Biol. 141:675–687.

Moritz, M., Y. Zheng, B.M. Alberts, and K. Oegema. 1998. Recruitment of the gamma-tubulin ring complex to Drosophila salt-stripped centrosome scaffolds. J Cell Biol. 142:775–786.

Murphy, S.M., A.M. Preble, U.K. Patel, K.L. O’Connell, D.P. Dias, M. Moritz, D. Agard, J.T. Stults, and T. Stearns. 2001. GCP5 and GCP6: two new members of the human gamma-tubulin complex. Mol Biol Cell. 12:3340–3352.

Murphy, S.M., L. Urbani, and T. Stearns. 1998. The mammalian gamma-tubulin complex contains homologues of the yeast spindle pole body components spc97p and spc98p. J Cell Biol. 141:663–674.

Nishimura, K., T. Fukagawa, H. Takisawa, T. Kakimoto, and M. Kanemaki. 2009. An auxin-based degron system for the rapid depletion of proteins in nonplant cells. Nat Methods. 6:917–922.

Oakley, B.R. 2023. The ring saga: looking back at the discovery of γ-tubulin and γ-tubulin ring complexes. Mol Biol Cell. 34:rt1.

Oegema, K., C. Wiese, O.C. Martin, R.A. Milligan, A. Iwamatsu, T.J. Mitchison, and Y. Zheng. 1999. Characterization of two related Drosophila gamma-tubulin complexes that differ in their ability to nucleate microtubules. J Cell Biol. 144:721–733.

Roostalu, J., and T. Surrey. 2017. Microtubule nucleation: beyond the template. Nat Rev Mol Cell Biol. 18:702–710.

Sundararajan, S., H. Park, S. Kawano, M. Johansson, B. Lama, T. Saito-Fujita, N. Saitoh, A. Arnaoutov, M. Dasso, Z. Wang, D. Keifenheim, D.J. Clarke, and Y. Azuma. 2023. Methylated histones on mitotic chromosomes promote topoisomerase IIalpha function for high fidelity chromosome segregation. iScience. 26:106743.

Tsuchiya, K., and G. Goshima. 2021. Microtubule-associated proteins promote microtubule generation in the absence of γ-tubulin in human colon cancer cells. J Cell Biol. 220.

Venkatram, S., J.J. Tasto, A. Feoktistova, J.L. Jennings, A.J. Link, and K.L. Gould. 2004. Identification and characterization of two novel proteins affecting fission yeast gamma-tubulin complex function. Mol Biol Cell. 15:2287–2301.

Verollet, C., N. Colombie, T. Daubon, H.M. Bourbon, M. Wright, and B. Raynaud-Messina. 2006. Drosophila melanogaster gamma-TuRC is dispensable for targeting gamma-tubulin to the centrosome and microtubule nucleation. J Cell Biol. 172:517–528.

Wieczorek, M., L. Urnavicius, S.C. Ti, K.R. Molloy, B.T. Chait, and T.M. Kapoor. 2020. Asymmetric Molecular Architecture of the Human gamma-Tubulin Ring Complex. Cell. 180:165–175 e116.

Woodruff, J.B., B. Ferreira Gomes, P.O. Widlund, J. Mahamid, A. Honigmann, and A.A. Hyman. 2017. The Centrosome Is a Selective Condensate that Nucleates Microtubules by Concentrating Tubulin. Cell. 169:1066–1077 e1010.

Wurtz, M., E. Zupa, E.S. Atorino, A. Neuner, A. Bohler, A.S. Rahadian, B.J.A. Vermeulen, G. Tonon, S. Eustermann, E. Schiebel, and S. Pfeffer. 2022. Modular assembly of the principal microtubule nucleator gamma-TuRC. Nat Commun. 13:473.

Xiong, Y., and B.R. Oakley. 2009. In vivo analysis of the functions of gamma-tubulin-complex proteins. J Cell Sci. 122:4218–4227.

Zeng, C. 2000. NuMA: a nuclear protein involved in mitotic centrosome function. Microsc Res Tech. 49:467–477.

Zheng, Y., M.L. Wong, B. Alberts, and T. Mitchison. 1995. Nucleation of microtubule assembly by a gamma-tubulin-containing ring complex. Nature. 378:578–583.

